# Common population codes produce extremely nonlinear neural manifolds

**DOI:** 10.1101/2022.09.27.509823

**Authors:** Anandita De, Rishidev Chaudhuri

## Abstract

Populations of neurons represent sensory, motor and cognitive variables via patterns of activity distributed across the population. The size of the population used to encode a variable is typically much greater than the dimension of the variable itself, and thus the corresponding neural population activity occupies lower-dimensional subsets of the full set of possible activity states. Given population activity data with such lower-dimensional structure, a fundamental question asks how close the low-dimensional data lies to a linear subspace. The linearity or non-linearity of the low-dimensional structure reflects important computational features of the encoding, such as robustness and generalizability. Moreover, identifying such linear structure underlies common data analysis methods such as Principal Component Analysis. Here we show that for data drawn from many common population codes the resulting point clouds and manifolds are exceedingly nonlinear, with the dimension of the best-fitting linear subspace growing at least exponentially with the true dimension of the data. Consequently, linear methods like Principal Component Analysis fail dramatically at identifying the true underlying structure, even in the limit of arbitrarily many data points and no noise.

## 1 Introduction

Neural coding is distributed and redundant, with large populations of neurons collectively encoding relevant variables. Geometric frameworks provide a natural setting within which to formulate and test theories of population coding, along with tools that allow population structure to be extracted from data [1–10]. In one particularly fruitful approach, data from a population of *N* neurons can be embedded in an *N* -dimensional space, with each axis corresponding to the activity of 1 neuron. The state of the population at each moment in time corresponds to a point in this *N* -dimensional space. Shared structure in the neural population code then corresponds to lower-dimensional shapes or, in the case of smooth responses, “manifolds” on which the data lies [11–20]. Computation can be understood in terms of trajectories on these low-dimensional manifolds [21–30].

Given some population data with such lower-dimensional structure, a fundamental question asks how close the data lies to a low-dimensional linear subspace or hyperplane (i.e., is the lower-dimensional structure near-linear?). This question is of theoretical interest because the linearity or non-linearity of the population data provides insight into the structure, robustness and generalizability of the encoding [9, 11, 31]. The linearity of data is also of great practical importance because methods that seek to fit a linear subspace to data, such as Principal Component Analysis (PCA) and Factor Analysis, are extremely widely used, whether to reveal structure in an unsupervised manner or as an initial data processing step before using regression and other supervised methods [1, 32]. Linear dimensionality reduction methods have a number of appealing features, including ease of interpretation, computational tractability, theoretical guarantees, and robustness. Moreover, linear methods are the foundation of a number of more advanced methods. For example, if a manifold is not well-fit by a linear subspace, a natural generalization is to seek a set of linear subspaces that combine to describe the manifold [33, 34]. On the other hand, using linear tools on highly non-linear manifolds will be misleading. Thus, understanding when a manifold is linear or near-linear provides insight both into the coding strategy being used by the corresponding brain region and determines the particular data analysis tools that can be used.

In this study, we examine the linearity of the manifolds generated by common population codes. We show that the resulting manifolds are exceptionally nonlinear. For example, consider a population of neurons with Gaussian tuning to a stimulus with *D* features—each neuron shows maximum response at some preferred stimulus value and the response decreases as a Gaussian function of the distance between the current stimulus and the maximally preferred stimulus value. Since there are *D* independent dimensions of variation, the neural population responses at any moment in time can be represented by a *D*-dimensional vector and the data is contained within a *D*-dimensional manifold. We prove, however, that a linear subspace that contains 80% (or any other fixed fraction) of the variance in these data must have dimension that grows exponentially with *D*. This dimension can be in the many thousands even for small values of *D*. Thus, methods that seek to fit a linear subspace to data will greatly overestimate the dimension of the true manifold, even in the limit of arbitrarily many data points and neurons.

## 2 Results

### 2.1 Setup

#### Low-dimensional population structure and neural manifolds

Given activity data from a population of *N* neurons over time, consider the population activity vector ***y***(*t*), whose *n*-th entry *y*_*n*_(*t*) is the activity (e.g., number of spikes fired or fluorescence signal) of neuron *n* in a time window of size *δt* centered at time *t*. This population activity vector can be seen as a point in an *N* -dimensional space, with each dimension of the space corresponding to the activity of one of the recorded neurons (see schematic in Fig. 1). If the population activity is measured at *T* time points *t*_1_, …, *t*_*T*_, then the recording yields *T* such population activity vectors. The population activity vectors together form the *N* × *T* data matrix *A*, where *A*_*ns*_ is the activity of neuron *n* in time bin *t*_*s*_.

**Figure 1.**
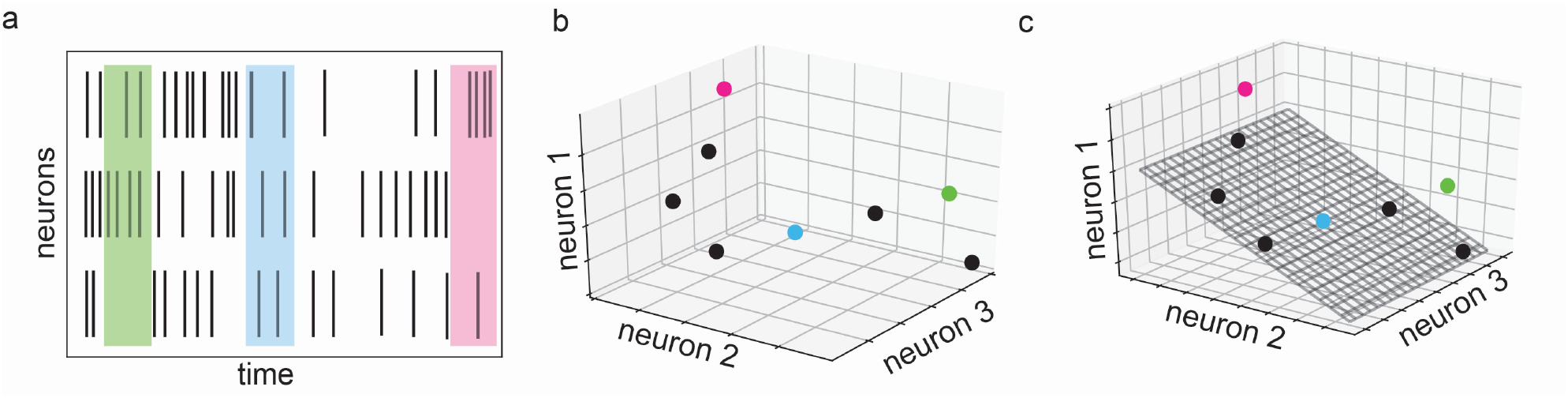
Schematic of low-dimensional structure in neural population data. (a) Spiking activity of 3 neurons over time. Shaded regions show three sample time bins, each used to compute an activity vector. (b) Activity represented as a collection of points in 3-dimensional space. Colored points correspond to shaded regions in panel (a). (c) Lower-dimensional linear structure in data, shown as a 2-dimensional plane chosen to capture as much variance in the data as possible. Scatter of points (e.g., pink and green points) off of plane reflects variance that is not captured.

The geometric picture corresponding to this collection of population activity vectors is a cloud of points in *N* -dimensional space, with each point corresponding to a moment in time (Fig. 1b). If the population of neurons shows structured activity, then the points will cluster around particular locations in the *N* -dimensional space or trace out particular shapes. These shapes provide ways to discover and to reason about the nature of the underlying representation or computation.

In particular, assume that the responses of the population of *N* neurons above are driven by some time-varying *D*-dimensional latent variable ***x***(*t*), meaning that ***y***(*t*) = ***F*** (***x***(*t*)) + ***ξ***(*t*) for some function ***F*** : ℝ^*D*^ → ℝ^*N*^, and where ***ξ***(*t*) is residual variance (e.g., noise). Here ***x***(*t*) could be an external stimulus, attentional or arousal or satiety state, motor plan, decision variable, internal estimate of location, or any combination of such and other variables. In what follows, we will adopt the terminology of Jazayeri & Ostojic [9] and refer to this stimulus or other population variable as the “latent variable”.

Note that, ignoring noise, the location of any population activity vector in the *N* -dimensional space can be specified by at most *D* coordinates (i.e., the values of the latent variable ***x***(*t*)) and thus the data point cloud lies in a *D*-dimensional space. We will refer to *D* as the “intrinsic” dimension of the data, following previous work [9]. Under some mild smoothness conditions on ***F*** and ***x***, the data lie on a *D*-dimensional manifold and we will thus refer to data “manifolds”, following common practice in the field (but the results do not require continuity and smoothness).

#### Populations with shared tuning curves

In the setting above, the response of the *n*th neuron is determined by *F*_*n*_, the *n*th component of ***F***. *F*_*n*_ thus captures the tuning of the *n*th neuron to the latent variable ***x***. In many neural populations, these tuning curves take a similar shape or functional form across neurons but differ in their preferred stimulus, width of selectivity, or other parameters (examples in Figs. 2a, 3a, 4a). Such shared tuning curve structure is common in topographically organized sensory regions [35–40] and in populations that show spatial tuning, such as place, grid and head direction cells [41–43]; it has also been found for more abstract quantities, such as among neurons tuned to numbers [44, 45] and to decision variables such as accumulated evidence [18, 46].

**Figure 2.**
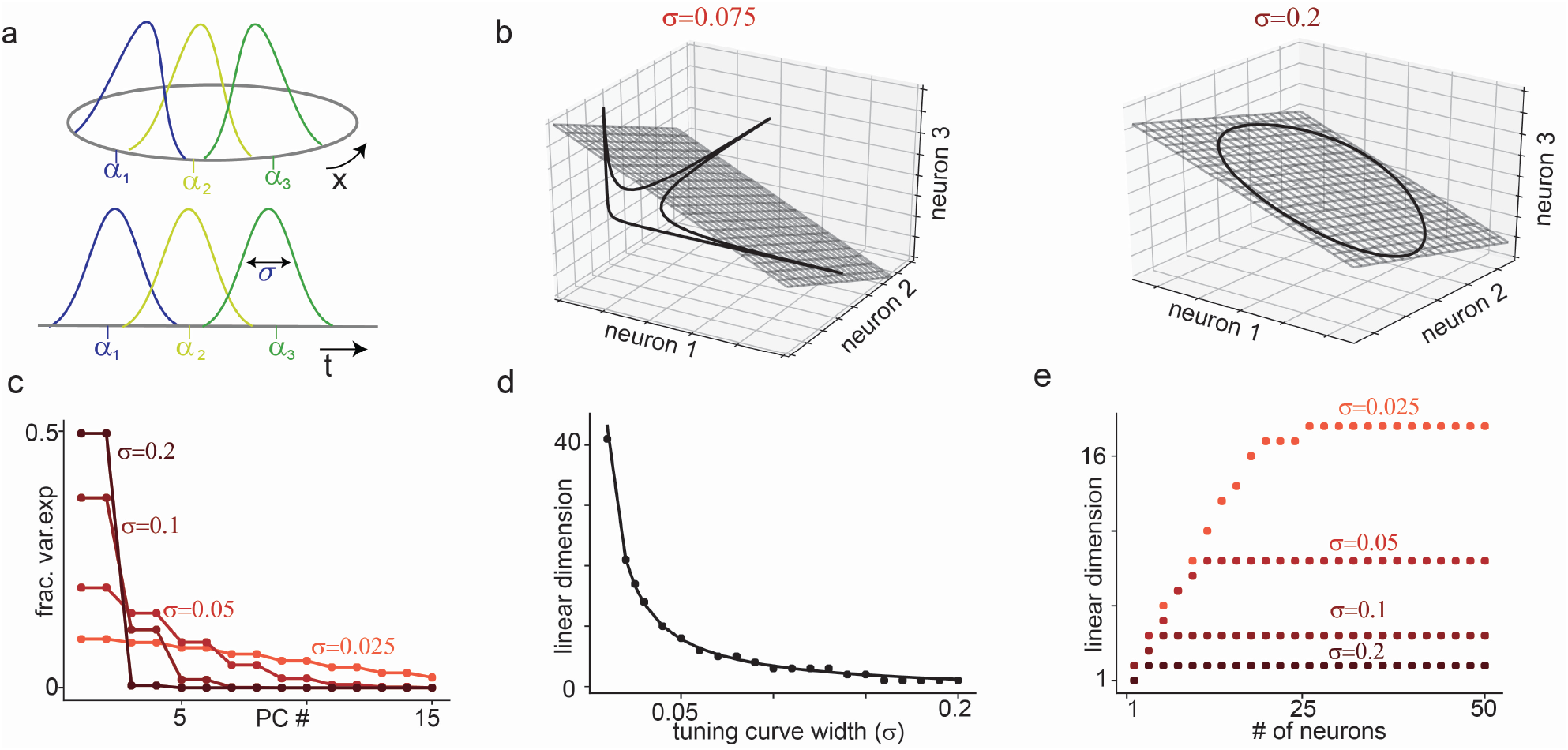
Translation-symmetric tuning to a one-dimensional variable and the inverse relationship between linear dimension and sparsity. (a) Gaussian tuning curves of 3 neurons encoding a circular (top) or non-circular (bottom) scalar stimulus variable. The non-circular variable example includes tuning to time, as in an epoch code. (b) Black line: Manifold formed by population activity of 3 neurons with Gaussian tuning to a 1-dimensional circular variable. Each axis shows the activity of 1 neuron. Gray: Best fitting 2D linear subspace (i.e., plane spanned by first two principal components). Left and right show an example of narrow (*σ* = 0.075) and broad (*σ* = 0.2) tuning respectively. For (c)–(e), results shown are for Gaussian tuning to a circular variable, with uniformly spaced tuning curve centers. Circles show numerical simulations and lines show theoretical predictions. (c) Fraction of variance explained by each principal component (equivalently, eigenvalues of covariance matrix) for a population of *N* = 50 neurons. Different curves show different tuning curve widths. (d) Linear dimension of neural data against tuning curve widths, showing that linear dimension grows as 1*/σ*. (e) Linear dimension against number of neurons in a population for each tuning curve width, showing initial linear growth before saturation at the predicted values shown in (d).

If tuning curves take a similar functional form, the activity of the *n*th neuron at time *t* can be modeled by *y*_*n*_(*t*) = *f* (***x***(*t*), ***α***_*n*_), with *n* = 1, …, *N*. Here *f* is a tuning curve function representing the shared shape of the tuning curve, ***α***_*n*_ is a vector of tuning curve parameters (e.g., preferred stimulus or tuning curve center of a sensory neuron, phase and period of a grid cell, preferred value of a decision variable, etc.), and, as before, ***x*** is the time-varying *D*-dimensional latent variable that underlies the population responses. Note that while such tuning curves were historically applied to describe the relationship between neural activity and an external variable, the latent variable can also be an internal cognitive variable [18, 46–51] or even an abstract statistical construct [13, 16, 52] that captures network interactions. For example, in a ring attractor network that encodes heading direction, the latent variable is the network’s estimate of heading direction, and the tuning of each neuron emerges from recurrent network interactions [42, 53, 54]. Thus the model is quite flexible and captures a variety of population codes.

#### Linearity of point clouds and manifolds

The most natural structure to seek in population data is linearity, corresponding to finding a lower-dimensional subspace (hyperplane) that contains the data (see schematic in Fig. 1c). If the dimension of this subspace is *L*, linear structure corresponds to finding *L* vectors ***v***_1_, *· · ·*, ***v***_*L*_ whose weighted sums account for the data. That is, any data point 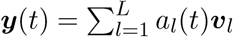, where *a*_*l*_(*t*) is the time-varying contribution of the *l*-th vector. Equivalently, the rank of the data matrix *A* is *L*.

In the presence of noise, data points will not lie exactly in a lower-dimensional linear subspace. Even in the absence of noise, a set of data points may not lie exactly in a linear subspace but might be close enough to be approximated by a linear subspace for practical purposes. Thus, it is typical to look for a linear subspace that captures most of the spread in the data while allowing for some scatter in the data around the subspace (Fig. 1c). Equivalently, one looks for *L* basis patterns that can approximately sum to any population activity vector 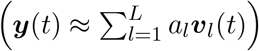 or for a rank *L* matrix *A*_*L*_ such that ||*A* − *A*_*L*_|| is small in some appropriate norm (usually 2-norm or Frobenius norm).

More precisely, we define the (1 − ϵ)-linear dimension *L*_1−*ϵ*_ of a matrix *A* to be the smallest *R* such that there exists a rank *R* matrix *A*_*R*_ for which 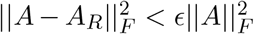 (this quantity is related to the *E*-rank of *A* [55]).

This definition of linear dimension corresponds to common practice in neural data analysis, where it is typical to perform PCA and estimate the dimension of data as the number of principal components required to explain some high fraction (i.e., 1 − ϵ in our notation) of the variance [1, 9, 56]. Thus, for example, what we call the 0.8-linear dimension is the number of principal components required to account for 80% of the variance in the data.

The best rank *R* approximation (or best fitting *R*-dimensional linear subspace) to the data matrix *A* is given by 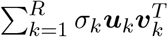, where *σ*_*k*_ is the *k*-th singular value of *A*, and ***u***_*k*_, ***v***_*k*_ are the *k*th left and right singular vectors respectively. In this case the remaining variance 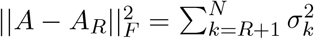. In other words, the matrix *A* has (1 − *ϵ*)-linear-dimension *L*_1−*ϵ*_ if

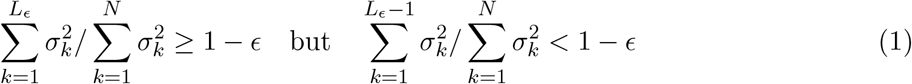

The singular values of *A* can also be calculated from the eigenvalues of the (non-mean-subtracted) covariance matrix *AA*^*T*^, which is the matrix of covariances between neurons averaged over time, or *A*^*T*^ *A*, which is the matrix of covariances between data points averaged over neurons. The *k*-th eigenvalue of each of these matrices is 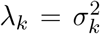 (for *k* ≤ *N*, assuming more time points than neurons).

Constructing such a low rank approximation to the data matrix (or, equivalently, fitting a linear subspace to the data point cloud) is the foundation of commonly used dimensionality reduction methods such as PCA and Factor Analysis. Moreover, a number of nonlinear dimensionality reduction techniques rely on approximating the data point cloud or manifold by a family of linear subspaces [33, 34, 57]. Such methods will be expected to perform well when the data point cloud or manifold is near-linear and poorly when the data manifold is highly non-linear.

#### Overview of approach

In this study, we consider a population of *N* neurons whose activity is driven by a *D*-dimensional real-valued latent variable ***x*** with firing rates given by tuning curve functions *f* (***x, α***_*n*_). Thus, the intrinsic dimension of neural activity is *D*. For several choices of tuning curve function we lower bound the (1−*ϵ*)-linear dimension of the neural activity (equivalently the number of principal components required to explain a (1 − *ϵ*) fraction of the data variance) and show that it is very large, growing at least exponentially with the intrinsic dimension *D*.

We assume that ***x*** takes on all possible values in a compact subset of ℝ^*D*^ and that *f* is continuous and one-to-one, so that in the absence of noise population responses lie on a *D*-dimensional manifold.

However, the approach can be naturally generalized to finding a linear subspace that contains point cloud data instead instead, and thus extends to cases like non-continuous values of the latent variable.

We consider firing rates and ignore noise so that the response of the *n*th neuron is given exactly by the mean firing rate, *y*_*n*_(*t*) = *f* (***x, α***_*n*_), and in this case the time window around *t* in which the rate is measured does not affect the results (as long as it is small on the timescale at which ***x*** changes). In the absence of noise, PCA and Factor Analysis are equivalent and our results thus apply to both methods (as well as to methods like Probabilistic PCA). Given that our results lower bound the linear dimension, including noise would simply strengthen our results by making neural activity more high-dimensional. Thus, our results reflect fundamental lower bounds on the dimensionality of neural activity rather than a lack of data and would not change if neural responses were averaged over multiple stimulus presentations.

We consider tuning curve functions *f* (***x, α***_*n*_) with certain symmetries and use these symmetries to exactly or approximately calculate the eigenvalues of the neuron-neuron covariance matrix (these eigenvalues are also the squared singular values of the data matrix and, when normalized, are the fractions of variance explained by the different principal components). We then count the number of eigenvalues needed to account for a (1 − ϵ) fraction of the variance in activity, for some small ϵ. Our results are not sensitive to the choice of *E* and in general apply when ϵ < 0.5.

To define the covariance matrix and calculate its eigenvalues, it will be convenient to first define the correlation profile function *c* between neurons with tuning parameters ***α***_***m***_ and ***α***_***n***_ to be *c*(***α***_***m***_, ***α***_***n***_) = 𝔼_***x***_ [*f* (***x***(*t*), ***α***_*m*_)*f* (***x***(*t*), ***α***_*n*_)], where the expectation is taken over the values of the latent variable ***x***. This is simply the (non-mean-subtracted) covariance between the neurons. For a population of *N* neurons, the *N* × *N* covariance matrix, *C*, has (*m, n*)th entry *C*_*mn*_ = *c*(***α***_***m***_, ***α***_***n***_).

Note that we consider the non-mean-subtracted covariance matrix except where otherwise indicated. Methods such as PCA often first subtract the mean from the data. In SI, Section 1 we show that if *L*_1−ϵ_ is the linear dimension of the non-mean-subtracted data, then *L*_1−ϵ_ − 1 is a lower bound on the linear dimension of the mean-subtracted data (this is a consequence of the Weyl inequalities relating the eigenvalues of perturbed matrices). Thus, our lower bounds on linear dimension for non-mean-subtracted data are easily transformed into lower bounds on mean-subtracted data, and in particular scaling arguments hold.

### 2.2 Translation-symmetric population codes

In many brain regions, responses to the latent variable are given by tuning curves whose shape is the same across neurons but with the tuning curve shifted or centered around a different region of latent variable space for each neuron [35–43]. Examples of such translation-symmetric tuning range from early sensory systems, such as orientation tuning in area V1 [37, 58], to cognitive systems, such as spatial tuning in hippocampal place cells and entorhinal grid cells [41, 43, 47, 48], epoch codes and hippocampal time cells [59], and tuning to abstract variables such as number [44, 45].

In this setting the response of neuron *n* at time *t* is determined by the difference between the current value of the latent variable, ***x***(*t*), and the neuron’s preferred value ***α***_***n***_ (note that for convenience we refer to the “center” or “preferred stimulus” of the tuning curve but the parameter ***α***_***n***_ more generally can simply index the shift of the tuning curve from an arbitrarily chosen reference tuning curve function as, for example, with the phase of a grid cell). If ***x*** ∈ℝ^*D*^ is the latent *D*-dimensional variable, then the tuning curve parameter ***α*** is also in ℝ^*D*^, and the tuning curve function *f* (***x, α***) = *g*(***x***−***α***) for some function *g* (equivalently, given some ***δ*** ∈ ℝ^*D*^, *f* (***x, α***) = *f* (***x***+***δ, α***+***δ***)). Different neurons have different preferred values, together tiling the space of possible values.

#### One-dimensional translation-symmetric populations and the sparsity-linear dimension uncertainty relation

First, consider a translation-symmetric population of *N* neurons where the encoded variable *x* and the tuning parameter for each neuron *α*_*n*_ are drawn from a one-dimensional circular space (i.e, S^1^), with points in the space parameterized by [0, 1), Fig. 2a (top). For example, *x* could be the angle of orientation of a bar, the direction of motion of a stimulus, or the head direction of a moving animal; correspondingly, *α*_*n*_ could be the center of the tuning curve of the *n*th neuron. If *x* evenly samples the space, then the correlation profile *c* for two neurons depends only on the the difference *α*_*m*_ − *α*_*n*_ between tuning parameters. We will slightly overload notation and write the correlation profile as *c*(*α*_*m*_, *α*_*n*_) = *c*(*δ*) where *δ* = *α*_*m*_ − *α*_*n*_ and the function *c* is periodic with period 1.

If the tuning parameters evenly tile the space, then the entries of the covariance matrix are

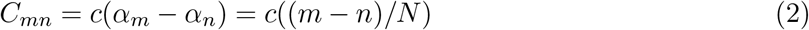

This covariance matrix *C* is circulant, meaning that each row is a shifted copy of the row above.

It is well known (and easy to show, see SI Section 2.1) that the eigenvalues of *C* are given by the Fourier transform of *c*, the function used to generate each row [4, 31, 60–62]. Thus, the *p*th eigenvalue of *C* is

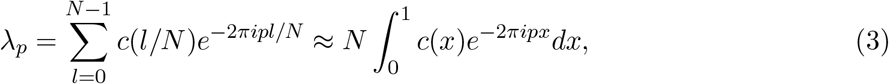

where 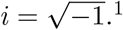

If the tuning curve centers are not evenly spaced, the corresponding matrix is no longer circulant, but the eigenvalues of the matrix *C* are still approximately given by the Fourier transform of *c*. If tuning curve centers are randomly sampled, this approximation converges rapidly as the number of neurons increases.

Similarly, consider a stimulus space that is one-dimensional but not circular, Fig. 2a (bottom). A natural example of this setting is tuning to time, as in an epoch code or the responses in a synfire chain. In this case the corresponding matrix is Toeplitz and the same eigenvalue relationship approximately holds [60, 63–66]. Thus, quite generally, the profile of eigenvalues is given by the Fourier transform of the correlation profile.

As a specific example, consider a population with translation-symmetric Gaussian tuning, which is a common model for tuning curves across multiple systems like orientation selective neurons in visual cortex V1 [67] and place cells in the hippocampus [41]. In this case the response of the *n*th neuron is modeled by

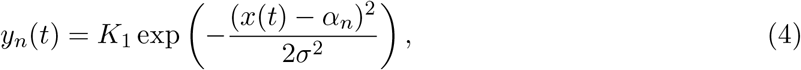

where *K*_1_ is the maximum firing rate and *σ* is the width of the tuning curve. The covariance between the *m*th and *n*th neurons is

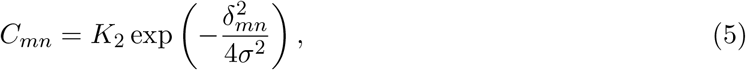

where *K*_2_ is a constant and *δ*_*mn*_ is the difference between the tuning curve centers (calculated accounting for the circular boundary conditions). The eigenvalues of the covariance matrix are given by

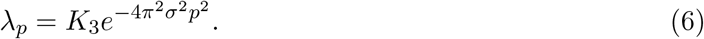

where *K*_3_ is a constant, as shown in Fig. 2C (see SI Section 2.3 and 2.4) for the calculation). Thus, the eigenvalue profile is Gaussian with variance inversely proportional to the width of the tuning curve.

The (1 − ϵ)-linear dimension is the smallest *L*_1− ϵ_ such that 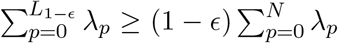. If *N* is not too small, these sums can be approximated by Gaussian integrals yielding the condition

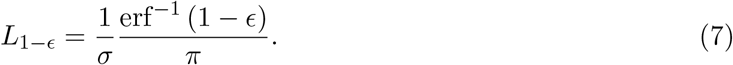

Thus, to explain a constant fraction of the variance, the linear dimension generically grows as 1*/σ*, where *σ* is the tuning curve width. In particular, *L*_0.95_ corresponds to a 95% confidence interval for a Gaussian distribution and is thus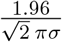.

More generally, so-called uncertainty principles relate the spread, sparsity, entropy or concentration of a function to that of its Fourier transform [68–71]. These principles imply that if the tuning curves (and hence the correlation profile) are sparse, fall off rapidly around their preferred values, or are concentrated on relatively small subsets of latent variable space, then *λ* will decay slowly with increasing *p* and have many significant non-zero entries. If *λ* decays slowly, then the number of eigenvalues needed to capture most of the variance will be high. Consequently, the linear dimension will be large and the manifold will be highly nonlinear.

Gaussian functions provide a lower bound for several of these uncertainty principles, and thus the 1/*σ* scaling relationship for Gaussians will provide a lower bound across a wide class of tuning curve shapes, in particular those with firing localized to some region of latent space (or the manifold). However, highly nonlinear manifolds are to be expected whenever the tuning curves (and hence the covariance profile) are concentrated on comparatively small subsets even if these tuning curves are not localized to a single interval or region.

We highlight two particularly useful uncertainty principles that apply in this more general setting. First, if the covariance profile *c* has *K* nonzero entries (i.e., it is *K*-sparse), then the eigenvalue profile has at least *N/K* nonzero entries [69]. Consequently, if *K* is small then there are a large number of nonzero eigenvalues. Second, if a fraction 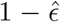 of the covariance profile is concentrated on a set *S* of size *K* (meaning that 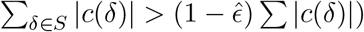, then the smallest set that contains 1 − *ϵ* of the eigenvalue mass has size at least 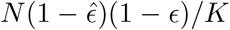 [69, 71]. Note that the size of this set is just the (1 − *ϵ*)-linear dimension and consequently the linear dimension again grows inversely with *K*. Sparse coding thus generically implies high linear dimension.

In practice, for systems with relatively broad tuning curves and for which the latent variable is low-dimensional, such as head direction cells [54, 72] or ventral hippocampal place cells [73, 74], the overestimate of intrinsic dimension by linear dimension may not be too large. However, in many systems, both sensory and cognitive, neurons respond to only a small fraction of possible values of the latent variable. For example, foveal V1 cells in the primate cover less than a degree of visual space [37], and rodent hippocampal place cells can cover under 1% of the area of large environments [74, 75]. Similarly, cells that are tuned to complex visual stimuli such as faces or other objects tend to show sparse responses [76, 77], thus covering only a small portion of stimulus space. In these settings, the manifold is likely to be highly nonlinear and linear dimensionality will greatly overestimate intrinsic dimensionality.

#### Multidimensional translation-symmetric tuning and exponential growth of linear dimension

As with the one-dimensional case, when a higher-dimensional variable ***x*** is encoded with translation-symmetric tuning curves (schematic in Fig. 3a), the covariance profile is also translation-symmetric and the eigenvalues of the covariance matrix (or singular values of the data matrix) are given by the Fourier transform of the covariance profile. Consequently, as in the 1D case, tuning curves that are sharper or concentrated on smaller sets will yield more slowly decaying eigenvalue profiles and hence higher linear dimension. However, the linear dimension will depend strongly on *D*, the intrinsic dimension of the latent variable. We first examine this interaction in the Gaussian case, before drawing general conclusions.

**Figure 3.**
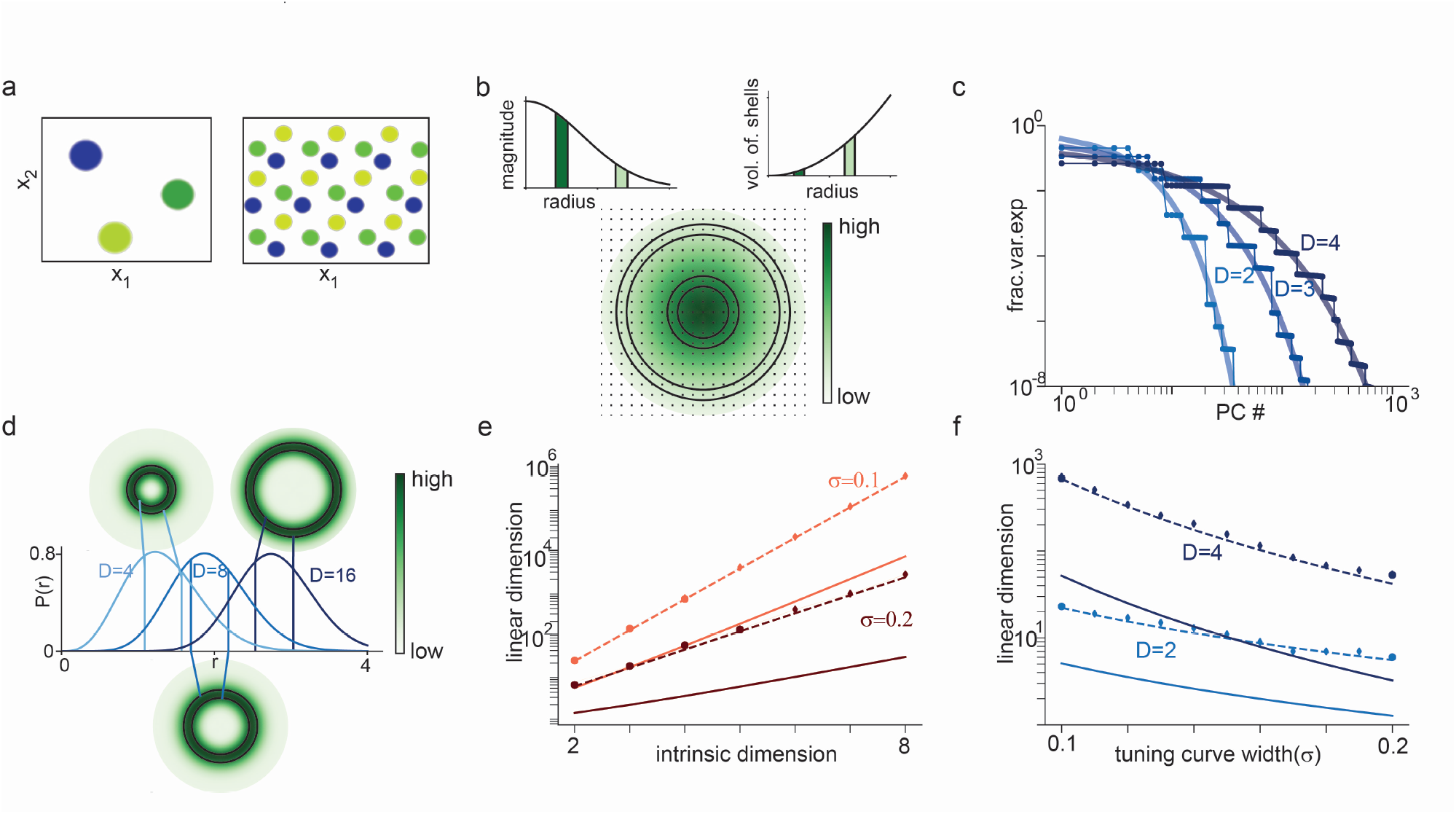
Translation-symmetric tuning to a multi-dimensional variable and exponential growth of linear dimension with intrinsic dimension. (a) Examples of 2d tuning curves, showing schematics of 3 different place cells with different tuning centers in a square arena (left) and 3 grid cells with the same spacing but different phases (right). (b) For Gaussian tuning curves, eigenvalues of the covariance matrix (variance along each PC) are values of a D-dimensional Gaussian at the lattice points of *D*-dimensional Fourier space. Each lattice point corresponds to one eigenvalue, and the colormap shows its value. Left inset: Decay of eigenvalues with distance from origin in Fourier space. Right: Number of eigenvalues contained in concentric shells of different radii. Circular shells on plot highlight two sets of eigenvalues, with corresponding magnitude and volume of shell shown as shaded region in insets. For a shell close to the origin the eigenvalues have large magnitude but there are fewer eigenvalues as a consequence of the smaller volume. Away from the origin the value of the eigenvalue is lower but there are more such eigenvalues. This tradeoff between eigenvalue magnitude and the number of eigenvalues of that magnitude explains the shape of the variance explained vs PC number curve. (c) Fraction of variance explained by each PC (or eigenvalues of covariance matrix) for *D*-dimensional Gaussian tuning curves and periodic boundary conditions along each dimension. Circles show numerical simulations, thin line represents prediction from Fourier transform of covariance matrix rows, and thicker lines represent theoretically-predicted smooth interpolation. (d) Total probability mass at radius *r* for a *D*-dimensional Gaussian (i.e., density function of chi distribution), shown for three different values of *D*. Circular insets show concentric shells colored by total probability mass at that radius. The bulk of the probability mass lies in a shell of radius 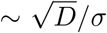. Thus, accounting for most of the variance requires considering all eigenvalues within a sphere of radius at least 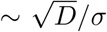. (e) Semi-log plot of linear dimension (*ϵ* = 0.05) vs. intrinsic dimension for Gaussian tuning curves with different widths. Circles show numerical results, solid lines show theoretical lower bound (applies whenever *ϵ* ≤ 0.5), and dashed lines show semi-analytic fit using chi distribution. (f) Semi-log plot of linear dimension vs. tuning curve width. Circles and lines as in panel (e).

Consider a population of *N* neurons with translation-symmetric Gaussian tuning to an underlying *D*-dimensional latent variable ***x*** that takes values within [0, 1] along each dimension. The tuning curve for the *n*th neuron is centered at ***α***_*n*_. By an appropriate choice of basis for ***x***, the covariance matrix of the Gaussian tuning curve can be assumed diagonal. For simplicity we assume that boundary conditions are circular and that ***x*** is scaled so that tuning curves have equal width *σ* < 1 in all directions. Thus, the response of the *n*-th neuron is

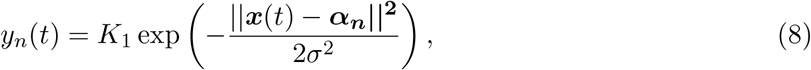

where *K*_1_ is the maximum firing rate. The corresponding correlation profile is also Gaussian. If tuning curve centers are equally spaced, then the covariance matrix has (*m, n*)th entry

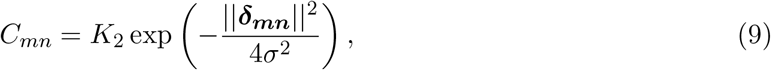

where *K*_2_ is a constant and ***δ***_***mn***_ is the difference between the tuning curve centers ***α***_***m***_ and ***α***_***n***_ (accounting for the circular boundary conditions). The eigenvalues are given by (SI Section 2.3 and 2.5)

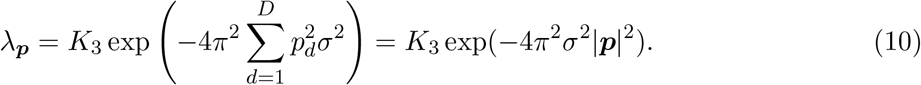

Here *K*_3_ is a constant. The eigenvalues are indexed by a *D*-dimensional vector ***p*** with *d*th entry *p*_*d*_ ∈ [−*N*_*d*_/2, *· · ·, N*_*d*_/2], where *N*_*d*_ is the number of tuning curve centers along dimension *d* (assumed the same for simplicity). Note that these eigenvalues are given by a multivariate Gaussian evaluated at the integer lattice points of a *D*-dimensional cube with side length *N*_*d*_, Fig. 3b.

The magnitude of an eigenvalue depends only on the magnitude of ***p*** and thus the eigenvalues vcan be ordered by smallest to largest distance from the origin in ***p***-space. There will be multiple eigenvalues with the same magnitude, corresponding to the same value of |***p***|. The number of eigenvalues of a given magnitude will increase with distance from the origin. Thus, moving away from the origin there will be more eigenvalues but of smaller magnitude (schematic in Fig. 3b).

When ordered by their magnitude, the eigenvalue profile thus shows a step-like shape, Fig. 3c. A smoothly interpolating function for the eigenvalue profile can be derived by noting that the eigenvalue profile is spherically symmetric in ***p***-space and thus the number of eigenvalues of a given magnitude depends on the number of lattice points at the corresponding radius. The number of lattice points can be interpolated by the volume of a sphere in *D*-dimensional space of radius *r*, which is given by

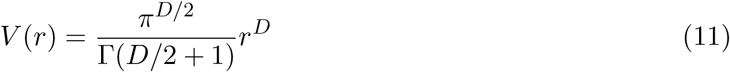

Thus, using Stirling’s approximation for the gamma function, the eigenvalues can be interpolated by the function

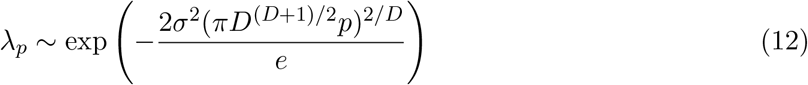

where the scalar *p* now indexes the eigenvalues from 1 to *N* (the total number of neurons) (also see [4] for a similar argument). The eigenvalues initially decay slowly and then rapidly, showing a transition between a power-law-like and an exponential regime.

To convert these eigenvalue profiles into an estimate of linear dimension, again note that the covariance profile is radially symmetric. Thus, we first seek the smallest radius *r* such that a fraction (1 − *ϵ*) of the total eigenvalue mass lies in a sphere of radius *r*, and then count the number of eigenvalues in that sphere. The scaling of this radius with *D* can be derived by observing that the probability mass of a *D*-dimensional Gaussian concentrates in a shell of radius 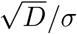 around the origin, as shown in Fig. 3d (we provide further details and calculate the radius more exactly using a *χ*-distribution in SI Section 2.5)

Thus, any sphere that captures a significant fraction of the probability mass must grow as 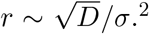 The number of eigenvalues (or lattice points) in a *D*-dimensional sphere of radius *r* grows as the volume, approximately as 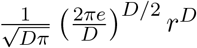. Consequently, as shown in Fig. 3e,f, the linear dimension grows as 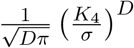, where *K*_4_ is a constant.

This scaling of linear dimension is extremely rapid, growing exponentially with intrinsic dimension. As a result, even low-dimensional Gaussian manifolds will have very high linear dimension. Moreover, this rapid growth means that even quite broad tuning curves will yield high linear dimension unless intrinsic dimension is extremely low. For example, as shown in Fig. 3e, a population of neurons with relatively broad Gaussian tuning to an 8 dimensional latent variable has linear dimension *L*_0.95_ > 6 × 10^5^.

The structure of the argument presented above is quite general, relying on the interaction between the decay of the eigenvalue magnitude with distance from the origin and the growth of volume with radius—while eigenvalues decay rapidly with distance, the growth of volume means that the radius of a sphere that captures some significant fraction of the total mass of eigenvalues must grow with *D*. The argument thus extends to other localized tuning curves even if non-Gaussian.

Exponential (or faster) scaling for localized tuning curves can be more generally derived from uncertainty principles. Analogous to the 1D setting, the results of [71] can be used to show that if a fraction 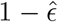 of the covariance profile is concentrated on a set *S* of size *K*, then the smallest set that contains 1 − *ϵ* of the eigenvalue mass (i.e., the (1 − *ϵ*)-linear dimension) has size at least 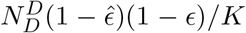 (where as before *N*_*D*_ is the number of tuning curve centers per dimension). For the case of Gaussian tuning, the size of the set containing 50% of the covariance profile can be upper bounded by the number of points in a sphere of radius 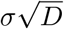, and this when combined with the uncertainty principle again yields exponential scaling (SI Section 2.5).

In the more realistic case where tuning curves are truly localized, meaning that each neuron’s tuning curve decays to some baseline value within a finite length (rather than, e.g., the small but infinite tails of a Gaussian function), most of the mass of the covariance profile is contained within a sphere of fixed radius, independent of dimension. In this setting, the linear dimension grows as 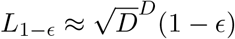 (see SI Section 2.6), and thus grows *supra-exponentially* with dimension.

Note that these results hold in the limit of arbitrarily many data points and recorded neurons.

In practice, the observed linear dimension will be limited by the number of neurons observed and the values of ***x*** sampled (or the task complexity [56]) but for these manifolds will continue to grow as more neurons and task variable values are measured.

### 2.3 Multiplicative tuning

We next consider tuning curve models where tuning to the latent variable can be written as a product over one-dimensional or lower-dimensional factors. That is, the tuning curve function is of the form

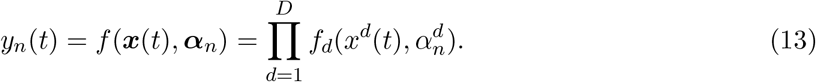

Here the superscript denotes the *d*-th component of a vector, and the *f*_*d*_’s are scalar functions. Note that in general the multiplicative factors can consist of groups of variables and not just one-dimensional factors, in which case the product would be taken over groups rather than single variables.

As an example of such tuning, if *f* is a multivariate Gaussian then, after an appropriate choice of basis for ***x***, *f* can be written as a product of one-dimensional Gaussians. As another example, common models of attention involve multiplicative gain modulation [78]. In this case the latent variable includes both the current stimulus value and the value of the attentional signal, and the response of a neuron can be written as a product of stimulus tuning and the response to the attentional signal (Fig. 4a (top)). For a third example, the spatio-temporal receptive fields of early visual cells are often decomposed as a product of the spatial part and the temporal part (Fig. 4a (bottom)) [79, 80]. Finally and more generally, multidimensional tuning curves that are not multiplicative may be able to be approximated by a product of lower-dimensional factors in a so-called “mean field” approximation.

**Figure 4.**
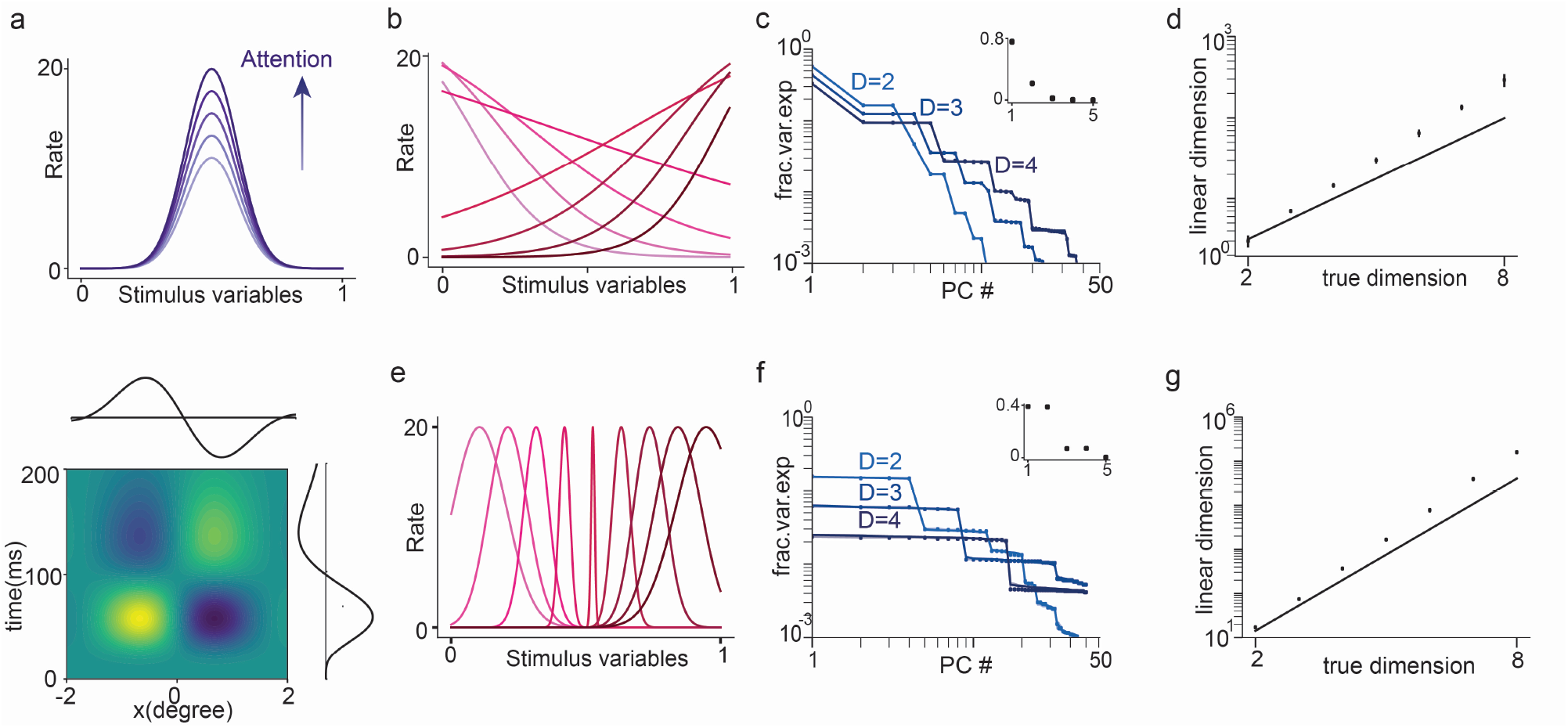
Multiplicative tuning and exponential growth of linear dimension with intrinsic dimension. (a) Schematics of common examples of multiplicative tuning. Top: Gain modulation of tuning to a sensory stimulus by attention. Bottom: Separable spatio-temporal receptive field of retinal ganglion cell as product of spatial tuning (horizontal) and temporal tuning (vertical). Panels (b)-(d) show results from a multiplicative tuning model where tuning along each dimension is sigmoidal. (b) Sample tuning along each dimension. Tuning curves are sigmoidal with slopes chosen uniformly in range [−5, 5]. (c) Fraction of variance explained vs PC number for the model shown in (b) for different values of intrinsic dimension (*D*). Circles show numerical simulations and lines show result from tensor product of 1D tuning curves. Inset shows the eigenvalues in the 1D case. (d) Linear dimension against intrinsic dimension for the data in (c). Circles show simulations and solid line shows theoretical lower bound of 2^*D*(*H*−0.05)^, where *H* is the entropy of the eigenvalue distribution shown in the inset of panel (c). Panels (e)-(g) show results from a multiplicative tuning model where tuning along each dimension is Gaussian. Gaussians are not translation-symmetric and the width of the Gaussian depends on position, with tuning sharpest at the center of the stimulus space (as in visual receptive fields). (e) Sample tuning along each dimension. (f), (g) As in (c), (d) but for the model shown in (e).

Let *m* and *n* be two neurons with parameter vectors ***α***_***m***_ and ***α***_***n***_. If the sampling of the encoded variable is uncorrelated across dimensions and boundary conditions are rectangular, then the covariance between these neurons can be written as

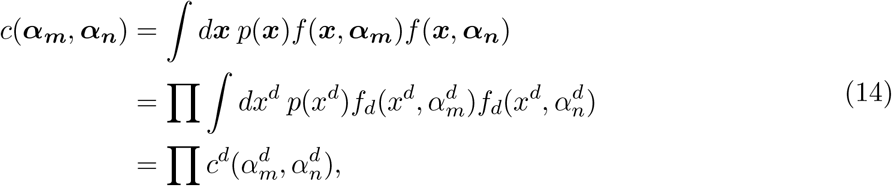

where *p*(***x***) = ∏_*d*_ *p*(*x*^*d*^) is the distribution of latent variable values and we have defined 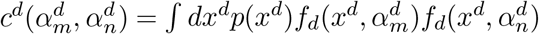.

If the tuning curve parameters are chosen to tile the space (note that tuning curves for different parameter values do not need to be shifted copies of each other, unlike in the previous section), then the data covariance matrix can be written as the tensor product of matrices corresponding to individual stimulus dimensions. Thus 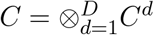, where ⊗ indicates the tensor product and the matrix *C*^*d*^ has entries 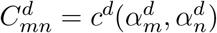 (see SI Section 3.1 for more details).

Now let 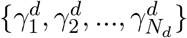 be the eigenvalues of *C*^d^, where *N*_*d*_ is the number of tuning curve parameters sampled along the *d*th dimension. The eigenvalues of *C* are all possible products of one eigenvalue from each *C*^*d*^ and so take the form 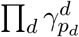, where 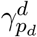 is the *p*_*d*_th eigenvalue of *C*^*d*^ and each *p*_*d*_ ranges over 0 to *N*_*d*_ − 1. In Fig. 4c,f we show examples of eigenvalues from two such architectures.

#### The linear dimension of multiplicative models grows at least exponentially with intrinsic dimension

Consider data from a *D*-dimensional multiplicative model where the tuning along each dimension has the same shape. Thus each component *C*^*d*^ has the same eigenvalues, which we denote {*γ*_1_, *γ*_2_, …, *γ*_*Nd*_ }.

To derive the asymptotic scaling of linear dimension with intrinsic dimension, we note that the eigenvalues of each component *C*^*d*^ (when appropriately normalized) can be interpreted as the outcome probabilities of a multinomial random variable. Moreover, the eigenvalues of the covariance matrix itself can be interpreted as the outcome probabilities of *D* independent draws from this distribution (see SI Section 3.2 for more details of the argument). Finding the smallest set of eigenvalues that sum to 1− *ϵ* of the total is thus equivalent to finding the smallest set of outcomes that accounts for 1 − *ϵ* of the total probability mass, sometimes called an *ϵ*-high-probability set [81]. The (1 − *ϵ*)-linear dimension is the size of this high-probability set.

The asymptotic equipartition property [81] guarantees that asymptotically the high-probability set contains 2^*DH*(***γ***)^ outcomes, where *H*(***γ***) = ∑_*p*_ *γ*_*p*_ log_2_ *γ*_*p*_ is the Shannon entropy of the eigenvalues of each component (normalized to sum to 1). Moreover, for any *E*^’^ > 0, 2^*D*(*H*(***γ***)− *ϵ*^*’* ^)^ is a lower bound on the size of the high-probability set for large enough *D*. This scaling is asymptotic, but in practice convergence is rapid and for the example models we consider the asymptotic lower bound captures the scaling even for small intrinsic dimension, as shown in Fig. 4d, g.

For a non-asymptotic lower bound on the linear dimension, assume that only two of the eigen-values for each factor are nonzero (and thus the overestimate of the intrinsic dimension of 1 by the linear dimension for each multiplicative factor is as small as possible while still being an overestimate). Up to an overall scaling constant (which we can ignore because we are considering eigenvalues as a fraction of the total sum), these eigenvalues can be written as {1 − *γ, γ*}, for some *γ* ≤ 0.5. The eigenvalues of *C* are again tensor products of [1 − *γ, γ*] taken *D* times. In descending order of magnitude, there is 1 eigenvalue of magnitude 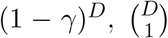 eigenvalues of magnitude (1 − *γ*)^*D*−1^*γ*, and so on, with 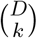 eigenvalues of magnitude (1 − *γ*)^*D*−*k*^*γ*^*k*^. As a consequence of our normalization, the sum of the eigenvalues 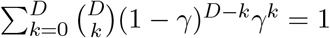.

Note that the eigenvalues are the outcome probabilities for a binomial random variable *X* distributed as Bin(*D, γ*) (i.e., *D* trials with success probability *γ*). Thus, the problem of finding *L*_1− *ϵ*_ is equivalent to first finding the smallest subset of outcomes of a binomial random variable that account for (1 − *ϵ*) of the probability and then counting the size of this subset.

Using standard bounds on sums of binomial coefficients yields a lower bound on the linear dimension for *ϵ* ≥ 0.5

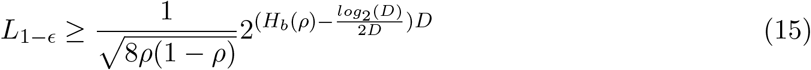

where *H*_*b*_(*ρ*) = −*ρ* log_2_ *ρ*−(1−*ρ*) log_2_(1−*ρ*) is the binary entropy function and *ρ* is lower bounded by *γ* − (1 + ln(2))*/D* (see SI Section 3.2 for full argument). Thus, except when *D* is small enough that 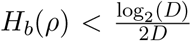 the lower bound grows exponentially with *D* (with the exponent asymptotically approaching *H*_*b*_(*γ*)*D*).

To summarize, for multiplicative tuning the linear dimension grows exponentially with intrinsic dimension, with a scaling constant that approaches the entropy of the eigenvalue distribution for a single factor.

## 3 Discussion

The relationship between intrinsic and linear (sometimes called “embedding”) dimension is emerging as an important means of understanding neural population coding, providing both insight into fundamental features of information encoding (such as generalizability and the progress of learning) and constraints on the statistical tools that can be used to analyze data [1, 9, 31]. It is widely appreciated that the point clouds and manifolds that emerge from neural population data are often nonlinear [9, 15, 18, 82], and previous work has in particular identified the sparsity of neural population responses as an important factor in this nonlinearity [4, 10, 17, 31]. The key contribution of the present study is to show that the nonlinearity is likely to be much greater than previously believed—for a number of common population codes, the linear dimension grows at least exponentially with the intrinsic dimension of the data. This exponential growth holds even if representations are not sparse, implying that even quite distributed population codes can have extremely high linear dimension. Consequently, dimensionality reduction methods that seek to fit a linear subspace to data, such as Principal Component Analysis and Factor Analysis, will dramatically overestimate the true dimension of data drawn from these population codes. Moreover, the exponential growth of linear dimension means that methods that fit data using a union of linear subspaces will require a large number of subspaces.

We have derived our analytical results for the cases of translation-symmetric and multiplicative tuning curves and find exponential or faster growth of linear dimension with intrinsic dimension in both settings. These results are likely to apply more generally to populations of neurons whose responses can be modeled as sparse or localized firing fields on some low-dimensional manifold, even if these firing fields take different shapes across neurons, as well as to populations whose tuning curves can be approximately written as a product of lower-dimensional factors in a so-called “mean-field” or separable approximation.

Our results on the high linear dimension of neural population data reflect the structure of the underlying manifolds or point clouds and do not reflect a lack of data or the presence of noise. In particular, they apply in the limit of large amounts of data and number of sampled neurons and in the absence of noise. For finite amounts of data, the observed linear dimension will be limited by the number of recorded neurons and the complexity of the task or experimental setting [56]. In the presence of noise, the observed linear dimension will be even higher than the noise-free calculations and thus the lower bounds will still hold. Finally, while we choose 90 − 95% variance explained as our criterion to define linear dimension for the figures, none of our results depend on this particular threshold, and exponential growth of linear dimension with intrinsic dimension is required to capture any non-vanishing fraction of variance.

Previous theoretical work measuring linear dimension has focused on the participation ratio [14, 56, 83–86]. If {*λ*_1_, …, *λ*_*N*_ } are the eigenvalues of the data covariance matrix, then the participation ratio (PR) is 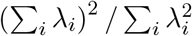. Thus, if the eigenvalue mass is concentrated on a few large eigenvalues the participation ratio is low. Notably, Recanatesi et al. show that for 2-dimensional Gaussian tuning, the PR increases inversely proportional to tuning curve width, much as we find [31]. We instead define linear dimension as the number of eigenvalues required to account for a certain fraction (i.e., 1 − *ϵ*) of total variance, as this is more easily interpretable in terms of data variance, and closely matches what is done in practice when using methods such as PCA [1, 9, 32, 82]. Depending on the particular eigenvalue distribution, the PR typically corresponds to the number of eigenvalues required to explain about 80% to 95% of the variance and is thus well-correlated with our definition of linear dimension [56]. Moreover, Wigderson & Wigderson [71] derive an uncertainty principle for localization as measured by the PR. In the context of our results, for translation-symmetric tuning this principle implies that the PR of the eigenvalue distribution must grow as the number of neurons divided by the spread of the covariance profile (similar to the growth of linear dimension with sparsity), where spread of the covariance profile is also measured using PR. Consequently, for translation-symmetric tuning our results will extend to linear dimension as measured by the PR. More generally, the PR has a number of nice mathematical properties and for these reasons was suggested as a more theoretically tractable alternative to fraction of variance explained when measuring dimensionality of neural population datasets [56]. Thus PR may provide a useful way to extend these results to other population codes.

The analyses presented here suggest that neural data from many brain regions should appear high-dimensional when viewed through linear dimensionality reduction methods, where “high-dimensional” is to be interpreted as large when compared to the number of encoded variables but still low-dimensional with respect to the number of neurons in a brain region. In accordance with our observations, recent data from large neural population data sets show high linear dimension [4, 87], and recordings from a number of low-dimensional systems appear distorted and significantly higher-dimensional than they actually are when viewed through linear methods as compared to nonlinear methods [15, 17, 18, 72].

Despite these observations, in many settings linear methods such as PCA and Factor Analysis have been successful at extracting structure from neural population data. What could explain this good performance?

One possible explanation is that the observed linear dimension is limited by task structure [56]. Gao and Ganguli show that the linear dimension (as measured by participation ratio) of neural data is upper-bounded by a measure of task complexity and argue that in common neuroscience tasks this measure is quite low. In many cases, the measure of task complexity they use grows exponentially with the number of task parameters. Thus, one possible test of the hypothesis that linear dimension is indeed limited by task structure is if observed linear dimension grows very rapidly as more task parameters are added. Another possible test is if linear dimension is higher in the case of naturalistic stimuli or during resting state activity when compared to more controlled task conditions, for which there is some evidence [88].

A second explanation for when PCA performs well is that the linearity or nonlinearity of neural point clouds and manifolds may differ sharply across different brain regions, reflecting differences in coding strategies, as suggested by recent work [9, 31]. Our results most naturally apply to coding in sensory systems, to circuits in the hippocampal formation that reflect spatial information, and to sparse combinatorial encoding of information in cognitive cortex. By contrast, if linear decodability is a good proxy for generalizability [6] then brain regions that strive to construct easily generalizable representations may show comparatively low linear dimension. Or, if the role of a brain region is to transform an initial condition into a particular dynamical pattern of activity, as suggested for motor cortex [89], then the data will be dominated by the linear dimension of the underlying dynamical system. This dynamical system may occupy a low-dimensional linear subspace because of constraints on learning and connectivity structure or the need for smoothness, controllability, and avoiding chaotic dynamics [3, 9, 12, 23]. As a third possibility, confining neural dynamics to low-dimensional linear subspaces that differ across tasks might enable efficient continual learning without interference [90]. Given these possiblities, as recently proposed the ratio of linear to intrinsic dimension might provide a useful signature of encoding strategies and task demands across brain regions and over the course of learning [9, 31]. Characterizing this ratio is an interesting direction for future work, and is increasingly tractable given advances in both large-scale recordings and nonlinear manifold learning algorithms.

Methods that seek to find the intrinsic dimension (*D*) of a nonlinear data manifold rather than using the dimension of a linear embedding (*L*) are an active and promising area of research [15, 17, 31, 34, 57, 82, 86, 87, 91–98]. Our formalism identifies a natural class of low-dimensional manifolds that should exist in neural data. These manifolds could be a useful and theoretically-tractable setting in which to evaluate existing and future methods that estimate intrinsic dimension, and thus to help identify promising candidates for real-world application. For example, a simple way to compare dimensionality estimation algorithms might be to apply them to simulated data generated from a population of neurons with Gaussian tuning to a *D*-dimensional latent variable, with added noise. The algorithms could then be compared on whether they successfully extract these highly nonlinear manifolds, how efficient they are at doing so in terms of both computation time and data, and how robust they are to noise in the data.

Finally, while our results suggest caution in the application of PCA and related linear methods, they raise the encouraging possibility that, at least in certain brain regions, low-dimensional structure in population responses may have been missed by linear analyses.

## 4 Methods

### Figure parameters and simulation details

Figure 2. For Figs. 2c,d, there are *N* = 50 neurons uniformly spaced in [0, 1] and *N*_*α*_ = 10^4^ values of the latent variable are drawn from a uniform distribution in [0, 1], so that *A* is a 50 × 10^4^ matrix.All neurons have Gaussian tuning curves with a fixed width *σ* and periodic boundary conditions. In Fig. 2c, circles are eigenvalues of the covariance matrix from simulations and lines are theoretical predictions from SI Eq. (22). In Fig. 2d, circles are *L*_0.95_ calculated from simulations, and the line is theoretical prediction in SI Eq. (24).

Figure 3. Circles in Fig. 3c,e,f are calculated from simulations with data generated from *D* dimensional Gaussian tuning curves with periodic boundary conditions along each dimension with the following parameters: *D* = 2, 10 neurons per dimension (100 total); *D* = 3, 10 neurons per dimension (1000 total); *D* = 4, 8 neurons per dimension (4096 total). Simulations used 10^4^ uniformly distributed values of the latent variable. In Fig. 3c, *σ* = 0.15. Thin lines are eigenvalues from theoretical prediction Eq. (10). Thick lines are smooth interpolations from Eq. (12). In Fig. 3e,f, diamonds are linear dimensions found numerically from tensor product of 1D eigenvalues (used to speed up computation as *D* gets larger). Dashed lines are semi-analytic approximation found from the chi-distribution. Solid lines are lower bounds in Eq. (11), and apply to any *L*_1− *ϵ*_ provided *ϵ* < 0.5.

Figure 4. For Fig. 3b-d, neurons along each dimension have sigmoidal response functions *f* (*x, α*) = 1/(1 + *e*^−*s*(*x*−*α*)^). For simulations, there are 8 neurons along each dimension with uniformly spaced *α* in [0, 1]. Slopes *s* are uniformly spaced in the range [−5, 5]. For Fig. 3e-g, neurons along each dimension have Gaussian tuning curves with varying widths, starting with the minimum value at the center of the range [0, 1] and increasing towards the ends. There are 8 neurons along each dimension. The minimum width is 0.05 and increases in steps of 0.05 to the maximum of 0.2. In Fig. 3c,f, circles are eigenvalues of covariance matrices calculated from *D* dimensional data and lines are tensor products of eigenvalues of the 1*D* covariance matrix, shown in the inset. In Fig. 3d,g, circles are *L*_0.9_ calculated from simulations and lines are asymptotic lower bounds 2^*D*(*H*−0.05)^ where *H* is the entropy of the normalised 1*D* eigenvalues shown in the inset of 3c,f.

## Acknowledgements

We thank Ila Fiete for helpful discussions and comments on this work, and John Murray for comments on the manuscript. This work was supported by funding from the Alfred P. Sloan Foundation under grant number FG-2021-16304. The authors were benefited by participating in the activities of the UC Davis TETRAPODS Institute of Data Science, which has been funded by NSF TRIPODS grant CCF-1934568.

## Supplementary Information

### 1 Linear dimension of mean subtracted data is lower bounded by the linear dimension of non-mean-subtracted data minus 1

We consider a population of *N* neurons with activities at time *t* given by ***y***(*t*) = [*y*_1_(*t*), *y*_2_(*t*), …, *y*_*N*_ (*t*)]. The mean of the population activity is ***μ*** = 𝔼_*t*_ [***y***(*t*)], where the expectation value is taken over time. The non-mean subtracted covariance matrix is given by *C* = 𝔼_*t*_ ***yy***^*T*^. And the mean-subtracted covariance matrix is given by *T* = 𝔼_*t*_[(***y*** − ***μ***)(***y*** − ***μ***)^*T*^]. Note that *C* = *T* + ***μμ***^*T*^.

The matrices *C, T* and ***μμ***^*T*^ are Hermitian and positive semi-definite. Moreover, ***μμ***^*T*^ is rank 1, with 1 non-zero eigenvalue ||***μ***||_2_ and remaining eigenvalues 0. Let the eigenvalues of these matrices be denoted as follows

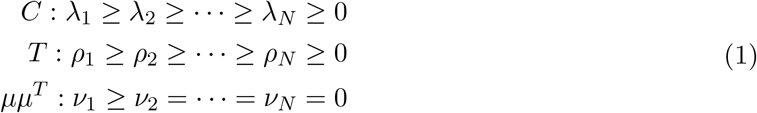

The Weyl inequality for Hermitian matrices yields

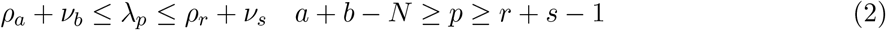

With *a* = *p, b* = *N, r* = *p* − 1, *s* = 2 we get

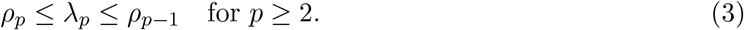

And with *a* = 1, *b* = *N, r* = 1, *s* = 1 we get

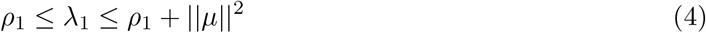

Thus we have the eigenvalue bounds

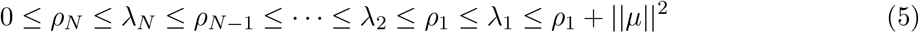

Now assume that the (1 − *ϵ*) linear dimension of the non-mean-subtracted system is *L* and that *L* is at least 2. Thus

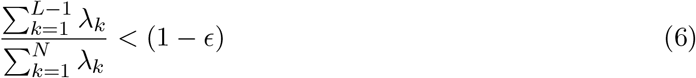

Define 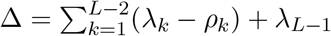. Note that 0 ≤ Δ ≤ *λ*_1_ by the above inequalities.

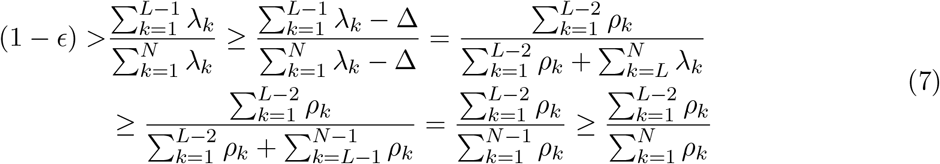

Thus 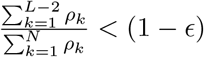.

Consequently if the (1 − *ϵ*)-linear dimension of *C* is *L*, the (1 − *ϵ*)-linear dimension of T is at least *L* − 1. In general, depending on ||*μ*||_2_ the linear dimension of *T* can be much higher than that of *C*. Thus, lower bounds on the linear dimension of the non-mean-subtracted case transfer simply to lower bounds on the linear dimension of the mean-subtracted case.

### 2 Translation-symmetric tuning curves

In this section, we derive the form of the covariance matrix for translation-symmetric tuning curves in Section 2.1 and show that the eigenvalues are given by the Fourier transform of the covariance profile in Section 2.2. We calculate the covariance matrix and its eigenvalues explicitly for translation-symmetric Gaussian tuning curves in Sections 2.3, 2.4, 2.5. We show that the linear dimension grows exponentially with *D* for Gaussian tuning curves in Section 2.5. Finally, we use an uncertainty principle relating functions and their Fourier transforms to show that the linear dimension for truly localized tuning curves grows supra-exponentially in Section 2.6.

#### 2.1 The covariance matrix of translation-symmetric data is translation-symmetric

We assume that the neural response depends on a *D*-dimensional latent variable, and thus the tuning centers, ***α***_***n***_ and the stimulus variables ***x***_*s*_ = ***x***(*t*_*s*_) are in ℝ^*D*^. The *ns*-th element of the data matrix is

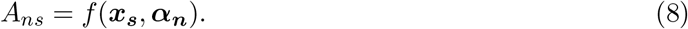

Here *f* is a translation-symmetric tuning curve so that *f* (***x***_***s***_ + ***δ, α***_***n***_ + ***δ***) = *f* (***x***_***s***_, ***α***_***n***_) = *g*(***x***_***s***_ − ***α***_***n***_) for some function *g*. We assume periodic boundary condition along each dimension with period 1 and thus ***x, α*** ∈ [0, 1]^*D*^. The latent variable is assumed to be uniformly distributed so that values are drawn uniformly from [0, 1]^*D*^.

The neuron-neuron covariance matrix is given by

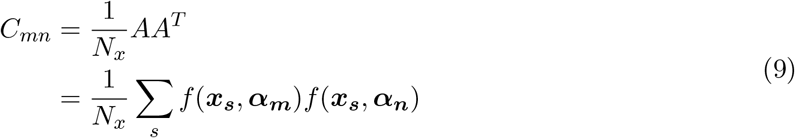

where *N*_*x*_ is the number of latent variable values. In the limit of large *N*_*x*_ this sample mean will converge to 𝔼 [*f* (***x, α***_***m***_)*f* (***x, α***_***n***_)]_***x***_, where the expectation is taken over the latent variable distribution. Thus the covariance matrix can also be written as

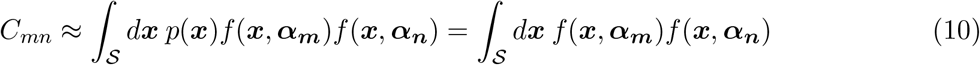

where *p*(***x***) is the the distribution over the latent variable, which we assume to be uniform, and the integral ∫_𝒮_*d****x*** = ∫_𝒮_ *dx*_1_ *· · · dx*_*D*_, is over all points ***x*** in the region 𝒮 = [0, 1]^*D*^. Since the tuning curves are translation-symmetric we have

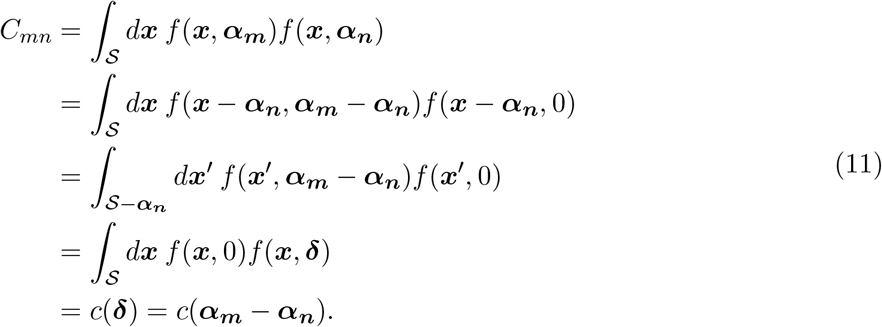

In going from the first to the second line, we have used the translation symmetry of the tuning curves (*f* (***x, α***) = *f* (***x*** − ***γ, α* − *γ***)); in going from the second line to the third line we have made a change of variable, ***x***^*t*^ = ***x*** − ***α***_***n***_ and defined ***δ*** = ***α***_***m***_ − ***α***_***n***_; and in going from the third line to fourth line we have used the periodic boundary conditions to shift the integral over 𝒮 − ***α***_***m***_ to 𝒮. *C* is a periodic function of the difference in tuning curve centers with period 1 along each dimension. Thus the covariance between two neurons just depends on the difference between their tuning curve centers.

In the one-dimensional case (*D* = 1), if we take the tuning curve centers to be uniformly spaced (i.e., *α*_*n*_ = *n/N* for *n* = 0, *· · · N* − 1), then the covariance matrix is circulant meaning that each row is a shifted version of the row above.

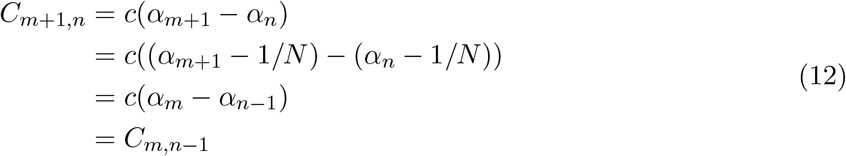

Note that this holds for *m, n* = 1, *· · · N*, since the periodic boundary conditions allow us to define *C*_*N* +1,*n*_ = *C*_1,*n*_ and *C*_*m*,0_ = *C*_*m,N*_.

Finally, note that the results approximately hold if the boundary conditions are hard rather than periodic, provided that the tuning curves are not too wide. In this case, the deviations from perfect translation-invariance caused by the boundary conditions will be mild and restricted to a small subset of neurons.

#### 2.2 Eigenvalues of translation-symmetric covariance matrices

It is well-known that matrices with translation-symmetric structure have eigenvalues that are given by the Fourier transform of the function that generates the matrix (i.e., *c*) [1, 2]. We briefly review those results here.

First, consider tuning curve centers that tile the latent space, forming the points of a lattice with *N*_*d*_ tuning curve centers along the *d*-th dimension. Thus, the spacing of tuning curve centers along the *d*-th dimension is 1*/N*_*d*_, and there are *N* = ∏_*d*_ *N*_*d*_ neurons in total, in a volume of 1^*D*^. The tuning curve centers are given by

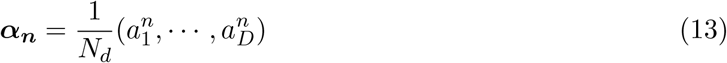

where each 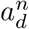 belongs to [0, 1, *· · ·, N*_*d*_ − 1].

The covariance matrix has entries *C*_*mn*_ = *c*(***α***_***m***_ − ***α***_*n*_) = *c*(***δ***). Note that because of the periodicity of *c* (as a consequence of the periodic boundary conditions), the vector ***δ*** can be considered to take the same set of possible values as the tuning curve centers.

Next, consider the set of *N* vectors ***k***_***p***_ given by

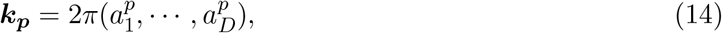

where as before each 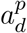 belongs to [0, 1, *· · ·, N*_*d*_ − 1]. Corresponding to each ***k***_***p***_, define the vector 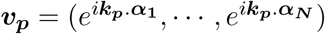. Multiplying this vector by the matrix *C* yields

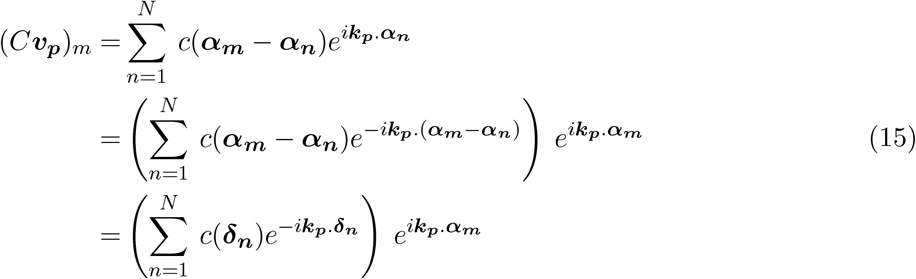

Here we have defined ***δ***_***n***_ = ***α***_***m***_ − ***α***_***n***_. As a consequence of the uniform spacing of tuning curve centers and periodic boundary conditions, the term in parentheses is a sum over all possible values of 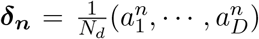, where as before each 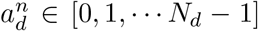. Thus, it is independent of both *m* and *n* and we can define 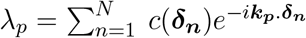. Consequently, ***v***_***p***_ is an eigenvector of *C* with eigenvalue *λ*_*p*_. Note that *λ*_*p*_ is the term with frequency *k*_*p*_ of the discrete Fourier transform of *c*.

In particular, for the one-dimensional case, *C* is a circulant matrix. In this case, we have tuning curve centers 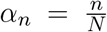, where *n* ∈ [0, *· · ·, N* − 1]. For each *p* with 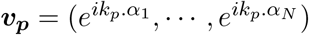 is an eigenvector with eigenvalue 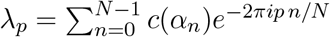.

Alternatively, if tuning curve centers are uniformly distributed through the latent space, either evenly spaced or randomly chosen, then in the large *N* limit we can consider the translation invariant kernel *c*(***α***_***m***_ − ***α***_***n***_), which is a continuous function of ***α***_***m***_, ***α***_***n***_ ∈ ℝ^*D*^. This is the continuous generalisation of the covariance matrix for translation-symmetric tuning curves derived above. The product of the matrix *C* with a vector can then be approximated by the convolution of the kernel with a function *f*_1_(***α***_***n***_),

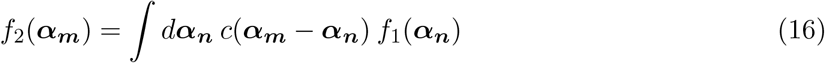

where ∫*d****α***_***n***_ means an integral over all *D* dimensional vectors ***α***_*n*_. *f* (***α***_*n*_) is an eigenfunction if

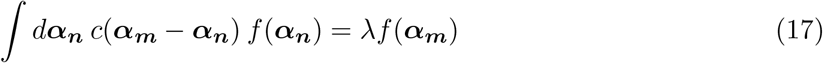

Consider 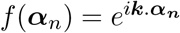 where ***k*** is an arbitrary *D* dimensional vector.

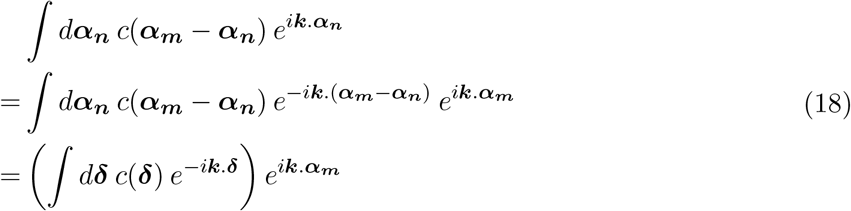

As before in going from line 2 to line 3, we have defined a new variable, ***α***_***m***_ − ***α***_***n***_ = ***δ*** and used the periodic boundary conditions to replace the integral over ***α***_***n***_ with an integral over ***δ***. The expression in parenthesis does not depend on ***α***_***m***_ or ***α***_***n***_ and is the Fourier transform of *c*(***δ***). So 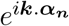 is an eigenfunction of *c*(***α***_***m***_ − ***α***_***n***_) with eigenvalue *d****δ*** *c*(***δ***) *e*^−*i****k*.*δ***^.

Finally, while these results are for uniformly distributed data with periodic boundary conditions, note that covariance matrices are symmetric and thus normal. Consequently, the effect of perturbations on the eigenvalue spectrum is as mild as possible, and the results should hold approximately for matrices that only approximately satisfy these conditions.

#### 2.3 Gaussian tuning curves show a Gaussian covariance profile

Consider the case of Gaussian tuning, where the tuning curve of the *n*-th neuron is given by

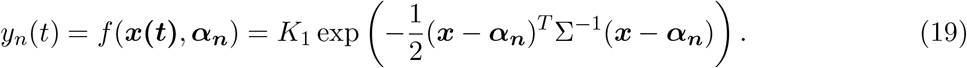

Here σ is the covariance matrix of the tuning curve.

To generate the neuron-neuron covariance matrix, we wish to calculate the covariance profile *c*(***δ***) = ∫ _𝒮_*d****x*** *f* (***x***, 0)*f* (***x, δ***), which is the covariance between pairs of neurons whose tuning curve centers are separated by ***δ***. Note that *c* is periodic, with period 1 in each direction. For notational convenience we shift the range of ***δ*** so that each component lies within [−1/2, 1/2] rather than [0, 1]. Thus,

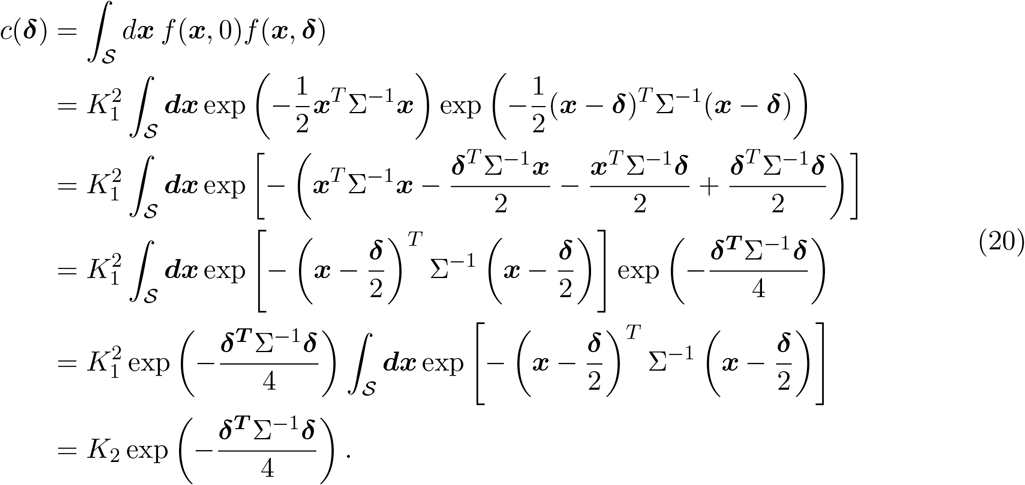

Here the constant 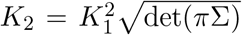. The last equality is approximate but holds when tuning curves are not too wide, meaning that no single neuron responds to the full range of latent variable values, so that integrating over the range of latent variable values is equivalent to integrating over the range of the tuning curve. Thus, neurons with Gaussian tuning curves have a Gaussian covariance profile with width twice that of the tuning curve.

#### 2.4 Variance explained and linear dimension for one-dimensional Gaussian tuning curves

In the one-dimensional case, the covariance profile is

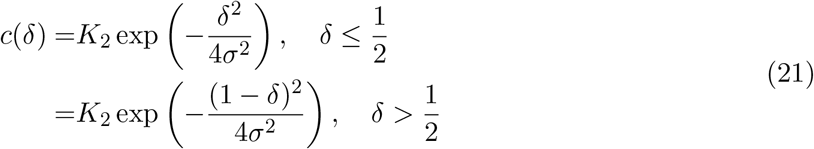

If tuning curve centers are evenly spaced, this covariance profile is sampled at *δ*_*n*_ = *n/N*, where *n* takes values in [0, …, *N* − 1].

As described in Section 2.2, the eigenvalues of the covariance matrix are given by the Fourier transform of this profile. For convenience we index the eigenvalues by *p* ranging from *l*(−*N* + 1)/2*J* to *lN*/2*J*. Ignoring the overall normalization constant *K*_2_, for each such *p* we have an eigenvalue

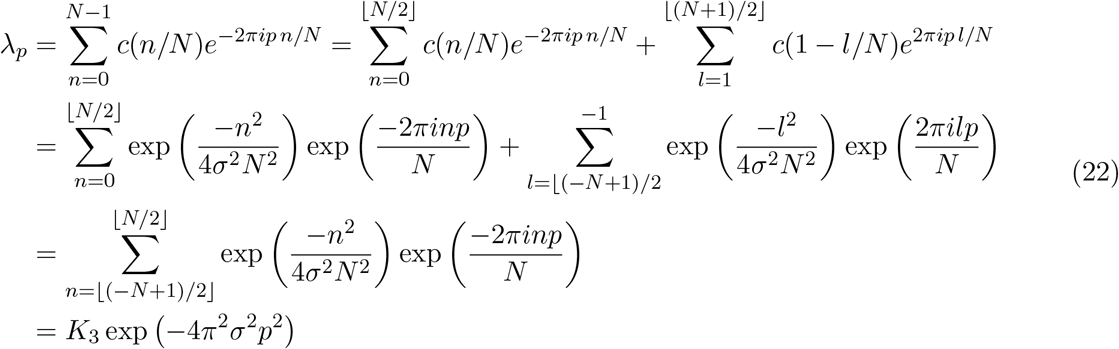

where *K*_3_ is a constant, and in the last line we have completed the square and noted that the sum over *n* is a constant. Thus the eigenvalues have a Gaussian profile with width 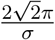. Also note that the eigenvalues decrease monotonically with the magnitude of *p* and that, except for *λ*_0_, they occur in pairs with *λ*_*p*_ = *λ*_−*p*_.

Since the eigenvalues occur in pairs, the (1 − *ϵ*)-linear dimension is the smallest *L*_1−*ϵ*_ such that

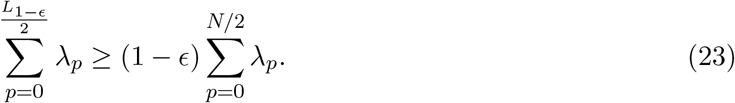

For large *N*, approximating both sides of this equation as an integral and canceling the common prefactor *K*_3_ yields

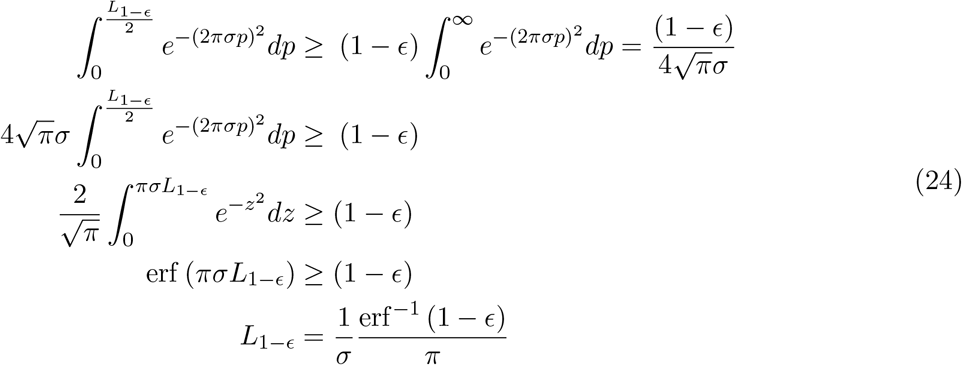

Thus, *L*_1−*ϵ*_ grows as 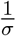, with a proportionality constant that depends on the fraction of variance explained. In particular, for 95% variance explained, we have 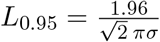.

#### 2.5 Variance explained and exponential scaling of linear dimension for multi-dimensional Gaussians

We next consider a population neurons with Gaussian tuning to an underlying *D*-dimensional latent variable. We assume that the tuning curve centers lie on a *D* dimensional grid with *N*_*d*_ equally spaced centers along each dimension. Therefore there are 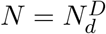 neurons.

For simplicity we also assume that the width of the tuning is the same along each direction so that the covariance matrix of the tuning curve σ is diagonal with elements *σ*^2^ along the diagonal.

Thus, as shown in Sec 2.2, the covariance between two neurons whose tuning curve centers are separated by ***δ*** is 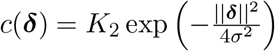, for 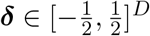, where *K*_2_ is an overall scaling constant. *c*(***δ***) is extended periodically outside this range.

As for the one-dimensional case and as discussed in Sec. 2.2, the eigenvalues of the neuron-neuron covariance matrix are given by the Fourier transform of *c*(***δ***). We can index the eigenvalues by a *D*-dimensional vector ***p*** with *d*th entry *p*_*d*_ ∈ [−*N*_*d*_/2, *· · ·, N*_*d*_/2] yielding

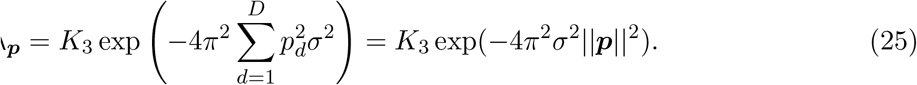

The magnitude of an eigenvalue is thus a function only of ||***p***||, the Euclidean distance from the origin in ***p***-space. As there are multiple lattice points with the same distance from the origin, there will be multiple eigenvalues with the same magnitude. Moreover, the number of eigenvalues at a given distance from the origin increases as the distance increases. Consequently, as the distance from the origin increases, there will be more eigenvalues but with smaller magnitude.

Overloading notation to define the *D*-dimensional continuous Gaussian function *λ* : ℝ^*D*^ → ℝ as *λ*(||***p***||) = *K*_3_ exp(−4*π*^2^*σ*^2^||***p***||_2_), observe that the eigenvalues are given by *λ*(||***p***||) sampled at the integer lattice points of a *D*-dimensional cube with side length *N*_*d*_.

To estimate the (1 − *ϵ*) linear dimension, note that summing up the *L*_1−*E*_ largest eigenvalues is equivalent to summing up all eigenvalues for which ||***p***|| ≤ *R* (i.e., eigenvalues corresponding to all lattice points with a ball of radius *R*), up to some edge effects resulting from the discreteness of the lattice points that will vanish when we take the continuous limit below. Thus we first compute the smallest radius *R* such that

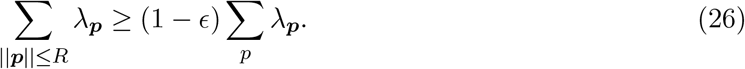

Approximating the sum by the integral of *λ*(***p***) and setting *K*_5_ = ; ∫_||***p***||_ *d****p****λ*(***p***), we seek *R* such that

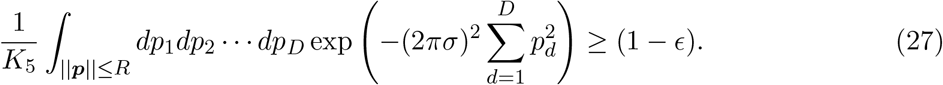

Converting to radial coordinates with 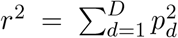 and then making a second change of coordinates 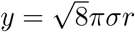 yields

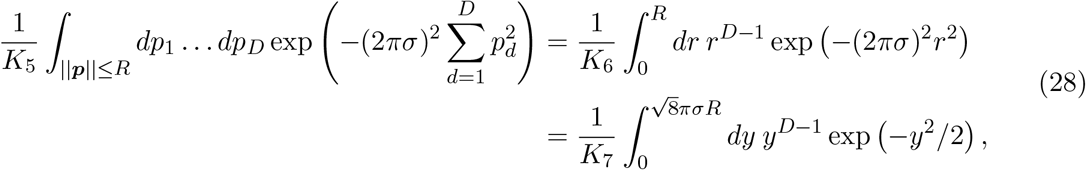

where 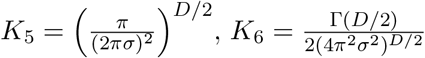 and *K* _7_ = 2^(*D*/2)−1^Γ(*D*/2).

Note that *λ* is the (unnormalized) probability density function of a multivariate Gaussian distribution and the argument of the integral above is the density function of a chi distribution with *D* degrees of freedom (appropriately normalized as a result of dividing by the integral across the entire range). Thus, defining the random variable *Z* distributed according to a chi distribution with *D* degrees of freedom, note that the integral itself corresponds to the CDF of *Z* and we seek the smallest *R* such that 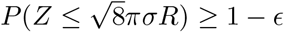.

If *ϵ <* 1/2 (i.e., we are capturing at least 50% of the variance in the data), then by definition *R* must be at le(ast the)median value of *Z*. Standard results on chi distributions then show that 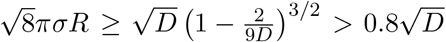, where the final inequality holds for *D* ≥ 2. Consequently, 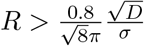.

Finally, to estimate the linear dimension itself we need to count the number of eigenvalues that lie within a sphere of radius *R*. Since the eigenvalues correspond to lattice points, the number of eigenvalues is approximately the volume of this *D*-dimensional sphere yielding

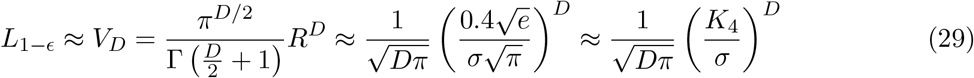

where *K*_4_ is a constant, and we have used Stirling’s approximation for the gamma function. Consequently the linear dimension grows exponentially with *D* as long as *σ* is not too large (note that the range along each dimension is normalized to [0, 1], so the threshold on *σ* effectively means that single tuning curves do not span the entire range of stimulus values).

#### 2.6 Supra-exponential growth of linear dimension for localized tuning curves from uncertainty principles

In this section we consider genuinely localized tuning curves, by which we mean tuning curves that are contained within some finite support rather than having small but infinite tails (i.e., unlike the Gaussian tuning curves in the previous section). We then use the uncertainty principles of Wigderson & Wigderson [3] to show that in this case linear dimension grows supra-exponentially with *D*.

We assume that the neurons have *N*_*D*_ equally spaced tuning curv e centers along each dimension in the hypercube [−1/2, 1/2]. Thus, there are a total of 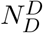 neurons in a volume of **1**. If tuning curves have finite support, then the tuning curve of a neuron is contained within some fixed radius *R*/2. We assume that an individual tuning curve does not cover more than half the space, so that *R*/2 < 1/4. Consequently, the covariance between two neurons is 0 unless the centers of their firing fields lie within a distance *R* < 1/2. Thus,

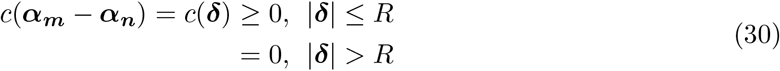

The support of the covariance profile *c* is thus upper-bounded by the number of points inside a sphere of radius *R*. Since there are 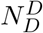 points in a volume of 1^*D*^, the support of the covariance profile *c* is,

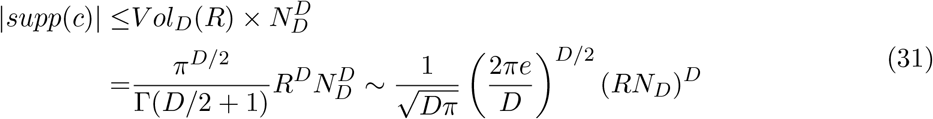

As before, the eigenvalues of the covariance profile are given by the Fourier transform of the covariance profile *c*. Let the eigenvalue profile be *λ*, with *ϵ* -support *supp* ^*ϵ*^ (*λ*). From the uncertainty principle for *ϵ* -support [3], we have

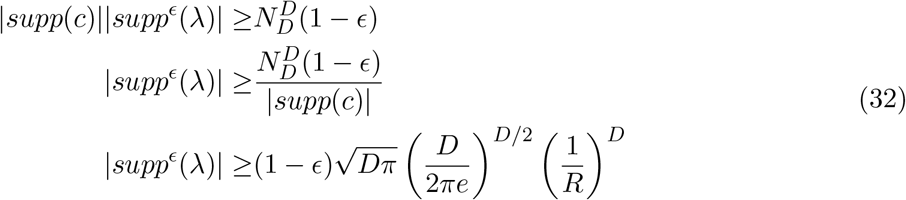

As before, the size of the *ϵ* -support of *λ* is the 1 − *ϵ* linear dimension. Thus, for translation-symmetric tuning curves which are localized within a fixed radius, the linear dimension grows faster than exponentially as 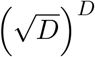.

### 3 Multiplicative tuning curves

In this section we show that for multiplicative tuning curves the D dimensional covariance matrix and its eigenvalues can be expressed as a tensor product of 1D covariance matrices and product of 1D eigenvalues respectively. We then find a lower bound on the linear dimension of data from such tuning curves using arguments from probability theory and information theory.

#### 3.1 Covariance matrix and eigenvalues of covariance matrix for multiplicative tuning curves

As in the main text we consider tuning curves of neurons which are a product of 1-dimensional tuning curves (but note that the argument extends to products over lower-dimensional factors even if these are not 1-dimensional).

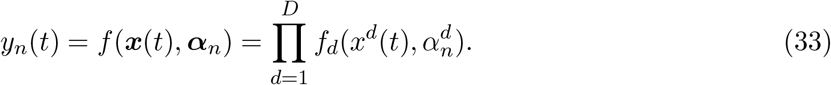

Assuming that the distribution of the latent variable along each dimension is independent (i.e., *p*(***x***) = ∏ _*d*_ *p*(*x*^*d*^)), the covariance profile between neurons with tuning parameters ***α***_***m***_ and ***α***_***n***_ is

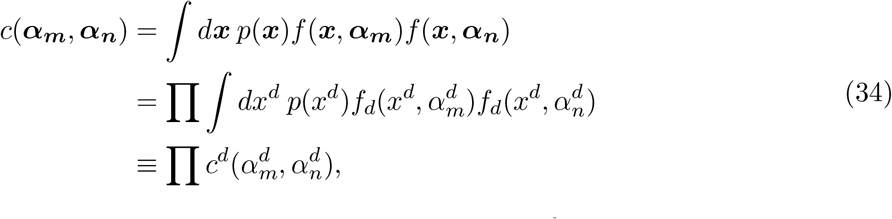

where the last line serves to define the individual covariance factors *c*^*d*^.

If the tuning curve centers are chosen to tile the space, then the covariance matrix *C* can be written as a tensor product

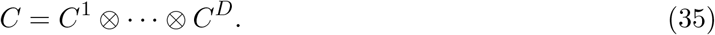

Here the factors 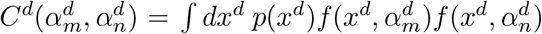, each correspond to a dimension of the latent variable.

Let 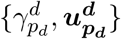 be the *p*_*d*_th eigenvalue-eigenvector pair for each *C*^d^. Note that

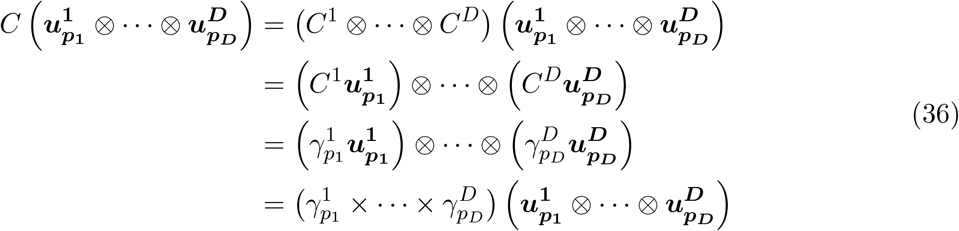

Consequently, 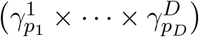 is an eigenvalue of *C*. All *N* eigenvalues of *C* can be constructed this way, as products of the eigenvalues of the factor *C*^*d*^. Consequently, the eigenvalues of *C* are given by all possible products of the eigenvalues of the individual factors.

#### 3.2 The linear dimension of multiplicative models grows exponentially with intrinsic dimension

As described in the main text, for simplicity, we will consider a multiplicative model where the tuning to each dimension has the same shape. Thus, each factor *C*^*d*^ is the same.

Let the eigenvalues of the covariance matrix for each factor be {*γ*_1_, …, *γ*_*ND*_ }. The *N* eigenvalues of the overall covariance matrix *C* are given by all products of the form ∏_*d*_ *γ*_*pd*_, where each *p*_*d*_ ∈ [1, …, *N*_*D*_]. To compute the linear dimension we reframe the problem as a problem in probability theory.

As we are considering fraction of variance explained, we can rescale {*γ*_1_, …, *γ*_*ND*_ } to sum to

1. Having made this normalization, consider a set of *D* independent random variables *Z*_*d*_, each distributed according to a multinomial distribution with outcome probabilities {*γ*_1_, …, *γ*_*ND*_ }. Each eigenvalue of *C* corresponds to the probability of some outcome of the string of random variables *Z*_1_ … *Z*_*D*_. Finding the smallest set of eigenvalues whose sum is at least 1 − *ϵ* (i.e., the *L*_1−*ϵ*_ linear dimension) is equivalent to finding the smallest set of outcomes whose probability is at least 1 − *ϵ*. This subset is often referred to as an *E*-high-probability set [4]. Standard results in information theory show that as *D* increases, the number of elements in this high-probability set approaches 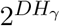, where *H*_*γ*_ = − Σ_*p*_ *γ*_*p*_ log_2_ *γ*_*p*_ is the Shannon entropy of the distribution {*γ*_1_, …, *γ*_*N*_ }. As each element in the high-probability set corresponds to an eigenvalue of the covariance matrix *C*, asymptotically *L*_1− *ϵ*_ grows as 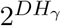, thus increasing exponentially with *D*.

While this is an asymptotic argument, exponential scaling holds for small *D* as well. For a non-asymptotic lower bound on the linear dimension of these models, consider the case where only two of the eigenvalues of the matrix *C*^*d*^ are nonzero. By normalizing to sum to 1, we can write these eigenvalues as 1 − *γ* and *γ*, for some *γ* ≤ 0.5. As for the multinomial case, the eigenvalues of *C* are the tensor product of [1 − *γ, γ*] taken *D* times, i.e. 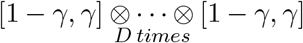. In descending order of magnitude, there is 1 eigenvalue of magnitude (1 *γ*)^*D*^, 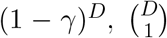 eigenvalues of magnitude (1 − *γ*)^*D*−1^*γ*, and so on, with 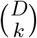 eigenvalues of magnitude (1 − *γ*)^*D*−*k*^*γ*^*k*^.

To lower bound the linear dimension, note that if *K* is such that

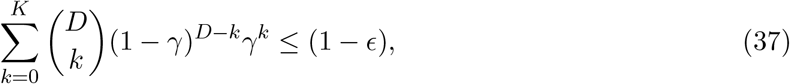

then the linear dimension 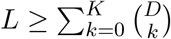. Thus, we will first find such a *K* and then use it to lowerbound the linear dimension *L*.

Consider a random variable *Z* distributed according to the binomial distribution with probability *γ*. That is, *X* ∼ *Bin*(*D, γ*). Note that 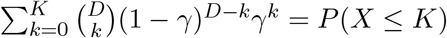. The median of this binomial distribution is at least ⌊*γD* ⌋. Thus if we choose *K* = ⌊*γD* ⌋ − 1 we have

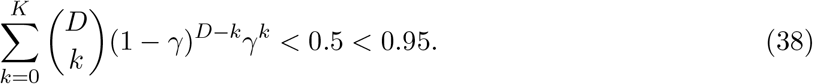

For simplicity, define *γ*, such that *γ*,*D* = ⌊*γD* ⌋ and note that we can rewrite *K* = *ρD*, where *ρ > γ* − 2*/D* (or, using results bounding the distance of the mean to the median o”£f a bin(om) ial, this can be improved to *ρ > γ* − (1 + ln(2))*/D*). We can now lower bound *L* as 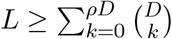.

A standard bound on the sum of binomial coefficients [4] yields

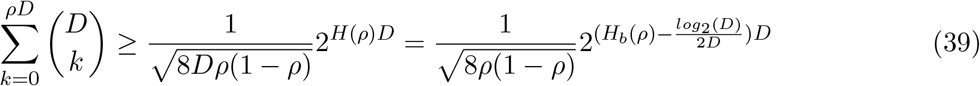

where *H*_*b*_(*ρ*) = −*ρ* log_2_ *ρ* − (1 − *ρ*) log_2_(1 − *ρ*) is the binary entropy function.

Thus, except when *D* is small enough that 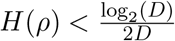 the lower bound grows exponentially with *D*, with the exponent asymptotically approaching *H*_*b*_(*γ*)*D*.

## Footnotes

1 Note that 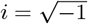 throughout this study and is not used as a matrix index.

2 Note that this problem is closely related to finding a 95% (or any other percent) confidence interval for a *D*-dimensional Gaussian). While an interval of width 2*σ* contains 95% of the probability mass in 1D, in higher dimensions an interval of any fixed width contains a shrinking fraction of the total probability mass. Thus the interval must grow with *D*.

